# ARID5B drives an inflammatory-to-destructive shift in pathologic fibroblast behavior

**DOI:** 10.64898/2026.08.04.742822

**Authors:** Angela E. Zou, Suppawat Kongthong, Gerald F.M. Watts, Cassandra L. Murphy, Madison L. Fairfield, Alisa A. Mueller, Michael B. Brenner

## Abstract

During inflammatory diseases such as rheumatoid arthritis, fibroblasts prominently drive chronic inflammation and the subsequent destruction of cartilage and bone. The mechanism by which an activated, inflammatory fibroblast acquires tissue destructive behaviors is unknown. Here, we describe ARID5B as a transcription factor that directs inflammatory fibroblasts to become migratory and invasive. Upon upregulation in inflammatory fibroblasts, ARID5B binds to histone editors and localizes to both inflammatory and invasive gene loci, epigenetically repressing pro-inflammatory genes while enhancing expression of pro-invasive genes. Likewise, fibroblast-specific ARID5B overexpression *in vivo* drives an inflammatory-to-erosive shift in arthritis pathology. Our findings highlight ARID5B as a maladaptive brake on inflammatory fibroblast activation that endows fibroblasts with pathologic invasive properties, thus mechanistically linking fibroblast-driven tissue inflammation to tissue damage. These insights into the regulation of inflammatory and invasive fibroblast pathology may inform successful therapeutic targeting of fibroblasts in inflammatory diseases.

## Introduction

Beyond structurally supporting tissues, healthy fibroblasts dynamically regulate local immune responses and actively remodel the tissue microenvironments in which they reside^1, 2, 3^. However, during chronic inflammatory and fibrotic diseases, pathologic fibroblasts drive persistent immune dysregulation and adverse tissue remodeling, including tissue destruction or fibrogenesis^1, 4^. These disease-associated fibroblast states contribute prominently to tissue damage and treatment resistance in conditions including cancer, inflammatory bowel disease (IBD), and rheumatoid arthritis (RA)^5, 6, 7, 8, 9, 10^.

RA is a systemic autoimmune disease marked by chronic inflammation of synovial joints and erosion of articular cartilage and bone. Mainstay RA therapies, including methotrexate and biologics such as anti-TNF and rituximab, primarily target the activities of inflammatory and autoreactive leukocytes^11, 12^. These drugs are efficacious, but leave a significant unmet need as only up to 40% of patients attain ACR70 treatment goals (≥70% reduction in disease activity) on any given therapy^12^ and up to 20-25% of patients remain refractory to multiple therapies^12, 13, 14^. Critically, recent work has revealed that fibroblasts persist as the most highly enriched cell population in the synovial tissues of patients with treatment-refractory RA^5, 9^. The Accelerating Medicines Partnership RA and Systemic Lupus Erythematosus (SLE) Network has further demonstrated at single-cell resolution that synovial tissues containing a predominance of fibroblasts are linked to resistance to anti-TNF and other DMARD therapies^9^. Thus, there is a need to better understand the mechanisms governing pathologic fibroblast activation that can be therapeutically exploited to foster disease remission.

During RA, inflammatory cytokines including TNFα, IL-17, IL1β, and IFNγ trigger the pathologic activation and expansion of synovial fibroblasts^15, 16^. Activated synovial fibroblasts in turn strongly drive tissue inflammation, becoming the primary producers of IL-6 in RA synovium and secreting high levels of CXCL8 (IL-8), leukemia inhibitory factor (LIF), and CXCL12 and other factors, which promote leukocyte recruitment and further amplify inflammatory fibroblast behavior^17, 18^. Moreover, activated fibroblasts within inflamed synovium acquire the ability to invade and destroy surrounding cartilage and bone, exhibiting increased migratory behaviors, inducing expression of matrix metalloproteinases (MMPs) and other extracellular matrix-degrading enzymes, and releasing RANKL to activate osteoclasts^19, 20, 21, 22^. Joint destruction is often irreversible and can cause permanent deformity and disability^11, 23^.

The major predictor of joint destruction in RA is preceding synovial inflammation^11, 24^. Longitudinal radiographic studies of RA joint pathology have shown that the degree of synovitis significantly determines subsequent progression of joint damage^25, 26, 27^, thus implicating synovial inflammation as the upstream driver of bone and cartilage erosion. However, the precise mechanisms by which the processes of tissue damage are triggered by local inflammation remains incompletely understood.

As fibroblasts play dual roles in perpetuating tissue inflammation and tissue invasion and destruction in RA, we hypothesized that they act as a critical cell type mediating the transition between inflammatory and invasive processes during disease. We specifically sought to examine whether a transcription factor (TF) that is activated in inflammatory fibroblasts can drive emergence of the invasive fibroblast state and mediates tissue destruction. TFs function as essential regulators of the phenotypic and functional programming of cells^28^. Recent studies have begun to identify TFs that orchestrate programs such as mechanosensing^29^, fibrosis^30^, and tissue remodeling by fibroblasts^31^. However, there is still a limited understanding of the TFs that govern fibroblast pathogenicity and effector function in RA and other inflammatory diseases.

Here, we find that the transcription factor AT-rich interaction domain 5B (ARID5B), previously linked to hematopoiesis and chondrogenesis^32, 33^ in other cell types, is selectively upregulated in inflammatory fibroblasts where it in turn promotes invasive fibroblast behavior. By complexing with histone editors and exerting opposing effects on the chromatin accessibility of target inflammatory versus invasive genes, ARID5B drives fibroblast invasiveness while epigenetically restraining their inflammatory activation. This ARID5B-controlled axis may serve as an attempted restraint on the pathologic inflammatory response that, instead of a one-way effect in reducing inflammation, acts through a mechanism of reciprocal chromatin changes that results in upregulation of aberrant invasive behavior. Together, these insights explain how the predominantly inflammatory tissue environment in RA drives the exuberant tissue damage response which is a major cause of disease progression and disability for patients with RA.

## Results

### Fibroblast-driven inflammation and invasion are positively correlated in human arthritis synovium

First, we sought to assess the relationship between inflammatory and invasive fibroblast activation in patients with arthritis. We analyzed transcriptional data from the Accelerating Medicines Partnership (AMP) RA/SLE Network Phase 2 study, which profiled 79,555 stromal cells from 81 OA and RA patient synovia stratified across ten subpopulation clusters^9^ (**Figure 1A**). For each patient, we scored synovial fibroblasts by their mean enrichment for a panel of genes known to define inflammatory fibroblast behavior and for a panel of genes known to define invasive and degradative fibroblast behaviors (**Table S1**). The inflammatory gene signature includes *IL6*, a key RA cytokine highly secreted by fibroblasts^17, 34^; *CXCL8* and *CXCL12*, leukocyte-recruiting chemokines produced by fibroblasts^17^; and *LIF*, a fibroblast-derived cytokine that amplifies proinflammatory fibroblast activation in an autocrine manner^18^. The invasive gene signature includes *MMP1*, *MMP13,* and *MMP14*, fibroblast-secreted collagenases that promote matrix degradation^19, 35, 36, 37^; *CDH11*, a mesenchymal cadherin that drives fibroblast invasiveness^21, 22^; and *TNFSF11* (RANKL), an osteoclast-activating factor secreted by fibroblasts^20, 38^. Across patients, mean enrichment of the inflammatory fibroblast signature positively correlated with mean enrichment of the invasive fibroblast signature (**Figure S1A**). This aligns with the concept that tissue inflammation increases erosive disease burden in RA^24, 25^, and suggests that inflammatory fibroblast activation may promote increased invasive behavior.

**Figure 1.**
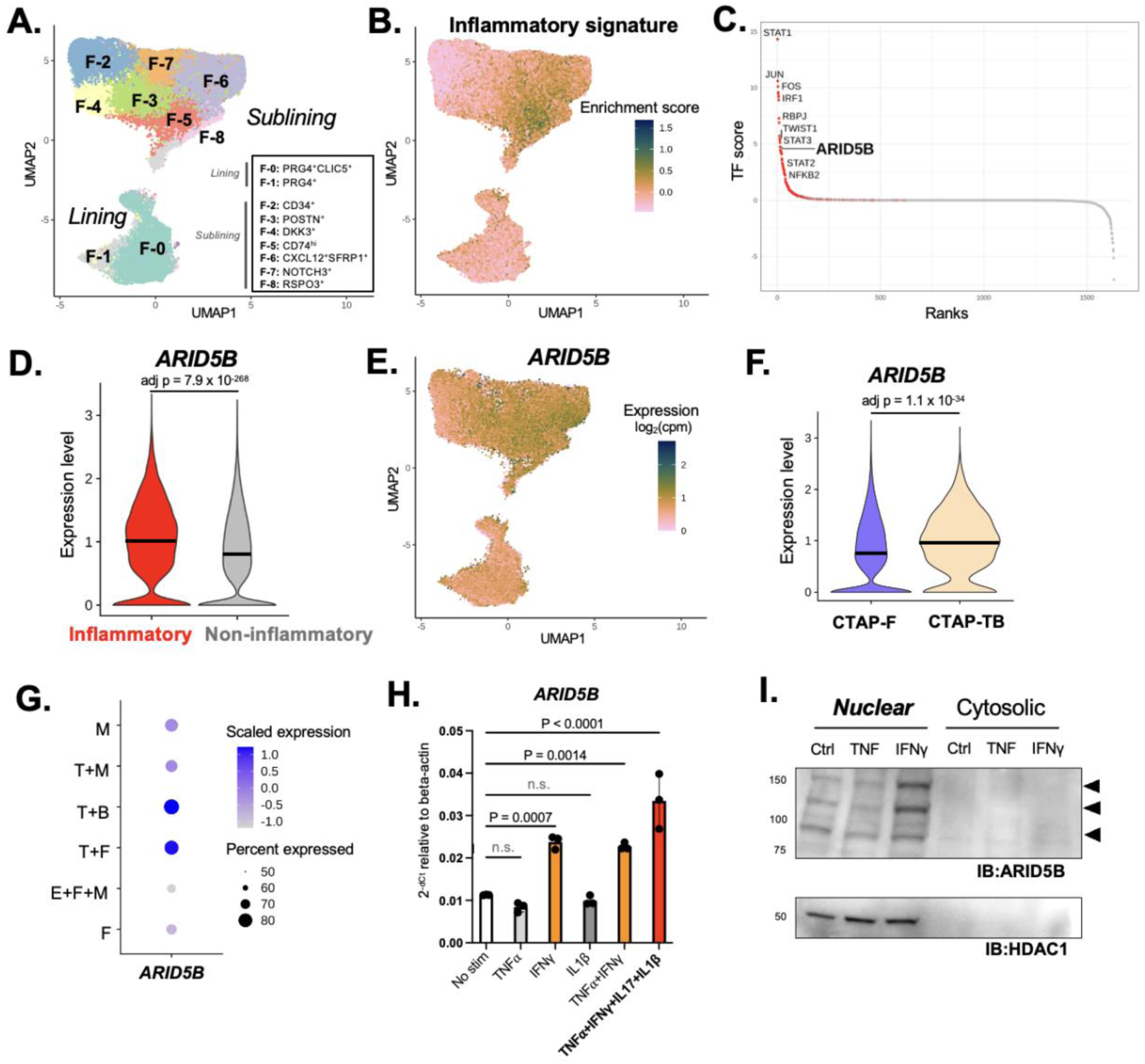
ARID5B is upregulated in inflammatory fibroblasts and induced by proinflammatory cytokines. **(A)** UMAP visualization of synovial fibroblast subpopulations identified in RA and OA patients from the AMP RA/SLE Network Phase 2 scRNA-seq dataset^9^. **(B)** Projection of inflammatory gene signature enrichment scores across synovial fibroblast subpopulations in UMAP space. **(C)** Dot plot depicting TF genes ranked by a scoring metric (TF score) corresponding to their differential expression in inflammatory versus noninflammatory synovial fibroblasts. Red points represent TFs significantly upregulated in inflammatory compared to noninflammatory fibroblasts (adjusted *p* < 0.05, from Wilcoxon rank-sum test with Bonferroni correction). **(D)** Violin plot of *ARID5B* expression in inflammatory versus noninflammatory fibroblasts. Median expression indicated; p value from Wilcoxon rank-sum test with Bonferroni correction. **(E)** Projection of *ARID5B* gene expression levels across synovial fibroblast subpopulations in UMAP space. **(F)** Violin plot of *ARID5B* expression in fibroblasts derived from CTAP-F versus CTAP-TB. Median expression indicated; p value from Wilcoxon rank-sum test with Bonferroni correction. **(G)** Dot plot of mean and proportion of *ARID5B* expression in fibroblasts derived from each RA CTAP. **(H)** RT-qPCR of *ARID5B* in primary synovial fibroblasts after stimulation with the indicated cytokines. Mean ± SD shown. P values from one-way ANOVA with Dunnett multiple comparisons test. **(I)** Immunoblot for ARID5B levels in nuclear and cytosolic synovial fibroblast lysates following stimulation with the indicated factors. ARID5B isoforms indicated by arrows. HDAC1 shown as nuclear lysate loading control.

### ARID5B is a transcription factor upregulated in inflammatory fibroblasts

Next, we examined inflammatory gene enrichment at the single-cell level to characterize the inflammatory fibroblast transcriptional state and identify upstream regulators driving activation of the invasive fibroblast state. The inflammatory gene module exhibits greatest enrichment among sublining *CD74^hi^* (F-5), *CXCL12^+^SFRP1^+^* (F-6), and *RSPO3^+^* (F-8) fibroblasts, which are known to express the highest levels of inflammatory factors such *IL6* and other cytokine response genes^9, 17^ (**Figure 1B, S1B**). Of note, *CD34^+^* fibroblasts (F-2) exhibit the lowest enrichment for the inflammatory gene module (**Figure S1B**), consistent with their proposed role as a homeostatic tissue-resident progenitor population^39^.

Given that inflammatory fibroblast activity is linked to increased fibroblast invasiveness, we next assessed whether a transcription factor (TF) upregulated in the inflammatory fibroblast state might govern the programming and activation of invasive fibroblasts. We performed a differential expression analysis of 1630 known TFs (as identified by Lambert et al.^28^) between fibroblasts most highly enriched for the inflammatory gene module and all other fibroblast populations. We subsequently scored TFs according to their degree of upregulation or downregulation and their expression levels in inflammatory versus noninflammatory fibroblasts (**Figure 1C**). Among the TFs most highly upregulated in inflammatory compared to noninflammatory fibroblasts are the AP-1 factors (*FOS*, *JUN)* and factors driving cytokine response pathways including interferon (*STAT1-3*, *IRF1*), and NF-κB signaling (*NFKB1, NFKB2*), all pathways established to promote synovial fibroblast inflammatory behavior^17, 40, 41^. Of interest, *ARID5B*, a TF with largely unknown functions in fibroblasts, was also significantly upregulated in inflammatory fibroblasts (**Figures 1C-E**), suggesting that it may directly regulate synovial fibroblast pathology.

Beyond its upregulation in inflammatory fibroblast states, we next assessed whether *ARID5B* is also heterogeneously expressed across fibroblasts derived from different RA patients. Across patients, mean enrichment of the inflammatory fibroblast signature positively correlated with mean fibroblast *ARID5B* expression, further supporting a direct relationship between fibroblast inflammation and *ARID5B* levels (**Figure S1C**). In the AMP RA/SLE study, patients with RA were also stratified into distinct Cell Type Abundance Phenotypes (CTAPs) based on the specific cell types most abundant within the synovial tissue^9^. Specifically, tissues designated as CTAP-TB contain a predominance of proinflammatory T and B cells, while tissues designated as CTAP-F are enriched in noninflammatory fibroblasts^9^. We find that ARID5B is significantly upregulated in fibroblasts derived from CTAP-TB compared to CTAP-F (**Figure 1F**). Moreover, across all defined CTAPs, *ARID5B* is more highly expressed in fibroblasts from other CTAPs rich in inflammatory leukocytes, including CTAP-M (myeloid cells), CTAP-TM (T cells and myeloid cells), CTAP-TF (T cells and fibroblasts) (**Figure 1G**), while being more lowly expressed in fibroblasts derived from CTAPs with lower proportions of leukocytes, including CTAP-F and CTAP-EFM (endothelial cells, fibroblasts, and myeloid cells) (**Figure S1C**).

### ARID5B is upregulated by proinflammatory cytokines

Given the increased levels of *ARID5B* in inflammatory fibroblast populations and in RA disease subgroups marked by active inflammation, we next sought to identify upstream regulators of ARID5B expression in fibroblasts. We stimulated primary synovial fibroblasts derived from RA patients with a panel of inflammatory cytokines implicated in RA, including TNFα, IFNγ, IL-17, and IL1β^15, 16^. Among these factors, IFNγ, both alone and in combination with TNFα, IL1β, and IL-17, most potently upregulated *ARID5B* gene expression in synovial fibroblasts (**Figure 1G**). Consistent with its function as a TF, ARID5B predominantly localizes to the synovial fibroblast nuclear compartment, and IFNγ stimulation upregulates ARID5B nuclear protein levels (**Figure 1H**). Thus, the proinflammatory cytokine milieu that promotes pathologic inflammatory fibroblast activation in RA also upregulates *ARID5B*.

We then assessed how regulation of ARID5B expression in fibroblasts may relate to RA pathogenesis. Genome-wide association studies have linked single nucleotide polymorphisms (SNPs) within the *ARID5B* locus to RA and SLE risk^42, 43, 44^. Through statistical fine-mapping, two SNPs (rs71508903 and rs7902146) located within the fourth intron of *ARID5B* were identified as being most likely causal in driving RA susceptibility^43^. To assess the potential relevance of these SNPs to *ARID5B* expression in fibroblasts, we performed ATAC-seq on two primary synovial fibroblast lines derived from RA patients and examined chromatin accessibility at the *ARID5B* gene locus. We observed that many SNPs within the *ARID5B* locus, including lead SNP rs71508903, overlap with ATAC-seq peaks corresponding to regions of open chromatin in synovial fibroblasts (**Figure S1D**), suggesting that these SNPs span active regulatory elements. Independently, our assessment of the *ARID5B* locus by GeneHancer^45^ found two regulatory elements overlapping with the SNP clusters which are predicted to modulate *ARID5B* expression levels (**Figures S1D**). Altogether, our findings indicate that ARID5B is upregulated by inflammatory stimuli and that variations in ARID5B gene regulation within fibroblasts may affect RA risk and disease presentation by altering the inflammatory or invasive effector functions assumed by activated fibroblasts.

### ARID5B binds the regulatory loci of genes encoding both invasive and inflammatory regulators in synovial fibroblasts

Given that ARID5B is upregulated in inflammatory synovial fibroblasts and is linked to RA risk and differences between RA subtypes, we sought to functionally characterize ARID5B by ascertaining its putative target genes in synovial fibroblasts. We performed CUT&RUN (cleavage under targets and release using nuclease) sequencing on synovial fibroblast lysates following ARID5B immunoprecipitation and enrichment for bound genomic regions using three distinct anti-ARID5B antibodies (**Figures 2A and S2A**). After peak calling, we prioritized our analysis to genes harboring regulatory regions bound by all three antibodies (**Figures 2B and S2B**). By Gene Ontology enrichment analysis^46^, we observed that these ARID5B target genes are significantly implicated in cellular processes regulating both inflammation and invasion (**Figure 2C**). Notably, within the *IL6* locus, both the gene promoter and proximal downstream regions exhibit peaks corresponding to ARID5B binding that are replicated among all three antibodies (**Figure 2D**), suggesting that ARID5B directly binds to and potentially regulates *IL6*. Apart from *IL6*, we observed ARID5B binding to the promoters and regulatory elements of other genes encoding inflammatory cytokines and chemokines (*CCL20*, *CXCL6*, *CXCL8*) and transducers of cytokine signaling (*JAK2*, *STAT4*) (**Figures 2E and S2B**). Additionally, ARID5B binds to key genes regulating cell motility and invasion, including adhesion molecules such as *CDH11*, Rho GTPase regulators that govern cytoskeleton organization (*ARHGAP*s and *ARHGEF*s), MMPs (*MMP13*), and mediators of epithelial-to-mesenchymal transition (EMT) including *TWIST1* and vimentin (*VIM*) (**Figure 2E**). These findings suggest that ARID5B is poised to directly regulate genes influencing both inflammatory and invasive functional states in synovial fibroblasts.

**Figure 2.**
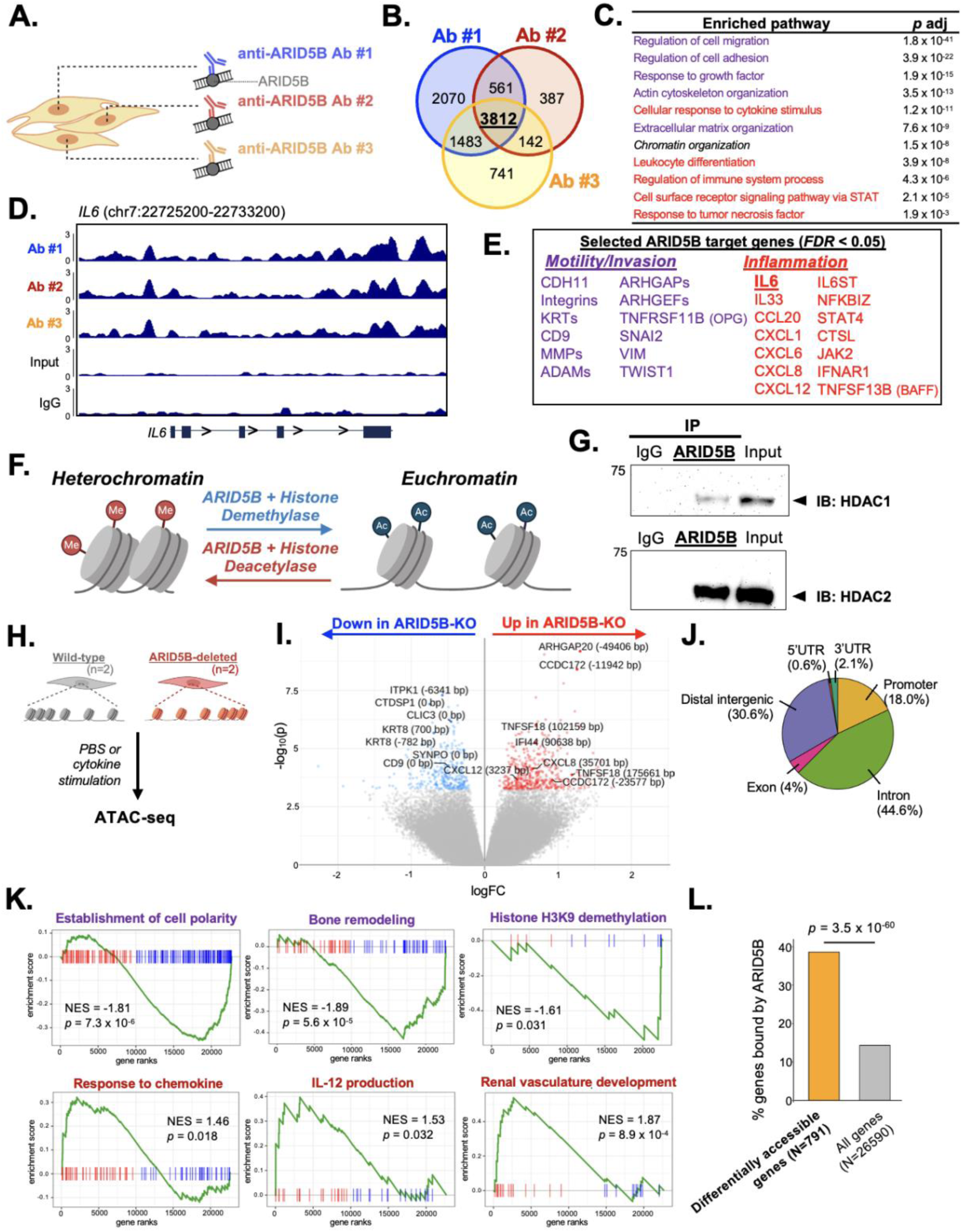
ARID5B directly binds invasive and inflammatory gene loci and reciprocally regulates their chromatin accessibility. **(A)** Experimental design for CUT&RUN sequencing of ARID5B in primary synovial fibroblasts. **(B)** Venn diagram enumerating the genes with regulatory regions captured by each anti-ARID5B antibody in CUT&RUN sequencing of synovial fibroblasts. **(C)** Selected Gene Ontology terms related to immune regulation and inflammation (red) and cell motility and invasion (purple) that are significantly enriched upon pathway analysis of ARID5B-bound genes. P values from Fisher’s Exact test with Benjamini-Hochberg correction. **(D)** Peak enrichment at the *IL6* gene locus for each ARID5B CUT&RUN library, compared to IgG isotype and input DNA negative controls. **(E)** Select genes bound by ARID5B in all three CUT&RUN libraries (*FDR* < 0.05 from Exact Poisson test with Benjamini-Hochberg correction). Genes colored in purple are related to migratory and invasive behavior and genes colored in red are related to immunity and inflammation. **(F)** Model of histone regulation by ARID5B-histone editor complexes. **(G)** Immunoblot for HDAC1 and HDAC2 following ARID5B pulldown from synovial fibroblast lysates, with nonspecific IgG pulldown as negative control and input lysate as positive control. **(H)** Experimental design for ATAC-seq of wild-type and ARID5B-deficient synovial fibroblasts. **(I)** Volcano plot depicting differentially accessible regions (DARs) in ARID5B-deficient fibroblasts versus controls. Red points indicate regions of significantly increased accessibility and blue points indicate regions of significantly decreased accessibility (*FDR* < 0.15). **(J)** Distribution of DARs between ARIDB-deficient versus control fibroblasts in relation to annotated genomic features. **(K)** Enrichment plots for selected pathway terms that are significantly negatively (purple) and positively (red) enriched among genes ranked by chromatin accessibility change between ARID5B-deficient versus control synovial fibroblasts (*p* < 0.05 by gene set enrichment analysis). **(L)** Enrichment of ARID5B binding at genes differentially accessible upon ARID5B deletion. P value from hypergeometric test.

### ARID5B deletion increases the chromatin accessibility of inflammatory gene loci and diminishes the accessibility of invasive gene loci

In leukocytes and chondrocytes, ARID5B functions as a chromatin-associated transcription factor, complexing with histone editors including demethylases and deacetylases (HDACs) to either promote or repress target gene expression^32, 47, 48^ (**Figure 2F**). Thus, we tested whether ARID5B also binds to histone editors in fibroblasts. Pull-down of ARID5B from primary fibroblasts followed by immunoblotting detected binding to both HDAC1 and HDAC2 (**Figures 2G and S2C**). *HDAC1* and *HDAC2* are broadly expressed across synovial fibroblast populations, including ARID5B-expressing fibroblasts (**Figures S2D and S2E**). This suggests that as in other cell types, ARID5B complexes with histone editors and functions as a chromatin regulator in fibroblasts, epigenetically controlling the expression of target genes.

We next sought to determine whether ARID5B alters chromatin states in synovial fibroblasts. We used CRISPR/Cas9 to delete ARID5B from primary human RA synovial fibroblasts (**Figure S2F**) and then performed ATAC-seq (assay for transposase-accessible chromatin sequencing) on both wild-type, non-targeting control (NTC) and ARID5B-deficient fibroblasts (**Figure 2H**). Following peak calling, ARID5B-deficient and control fibroblasts separate by principal components analysis (PCA) (**Figure S2G**). We detected 791 differentially accessible regions (DARs) between ARID5B-deficient versus control fibroblasts (*FDR* < 0.15, **Figure 2I**), predominantly localized to promoter, intronic, and distal intergenic regulatory regions (**Figure 2J**). We then performed gene set enrichment analyses (GSEA)^49^ on all genes ranked according to change in accessibility following ARID5B deletion. In ARID5B-deficient fibroblasts compared to control fibroblasts, we observed decreased chromatin accessibility among genes involved in cell polarity and bone remodeling pathways, both associated with fibroblast motility and tissue degradative behavior (**Figure 2K**). We also observed that ARID5B deletion reduced the chromatin accessibility of genes mediating histone demethylation processes (**Figure 2K**), consistent with the proposed histone regulatory function of ARID5B. In contrast, we observed enhanced chromatin accessibility among genes involved in immune activation, cytokine signaling, and angiogenesis-related pathways, suggestive of inflammatory fibroblast activation (**Figure 2K**).

Finally, we assessed whether genes regulated by ARID5B at the chromatin accessibility level correspond to genes directly bound by ARID5B at one or more regulatory loci. Indeed, among genes exhibiting differential chromatin accessibility following ARID5B deletion, we observe a significant enrichment of genes identified to be bound by ARID5B from CUT&RUN sequencing (*p* = 3.5 x 10^-60^, **Figure 2L**). Collectively, these findings show that ARID5B, in association with histone editors, binds to and alters the chromatin accessibility of target inflammatory and invasive gene loci. Specifically, when ARID5B is induced by inflammatory stimuli, it can in turn diminish the accessibility of genes mediating the fibroblast inflammatory response and enhance accessibility of genes mediating invasive fibroblast behavior.

### ARID5B suppresses inflammatory fibroblast activation

Given that ARID5B binds inflammatory genes and that its loss increases inflammatory gene accessibility, we next functionally tested the effects of ARID5B on the fibroblast inflammatory state. We generated wild-type control (NTC) and ARID5B-deficient primary synovial fibroblasts and compared their responses to stimulation with the cytokine combination of TNFα, IFNγ, IL-17, and IL1β which was found to most highly induce *ARID5B* expression (**Figure 1G**). Upon cytokine treatment, we observed a significant increase in both IL-6 and CXCL8 secretion (**Figure 3A**) and gene expression (**Figure 3B**) in ARID5B-deficient synovial fibroblasts compared to control. Likewise, monostimulation with TNFα also enhanced expression of *IL6* and *CXCL8* in ARID5B-deficient fibroblasts over control (**Figure S3A**).

**Figure 3.**
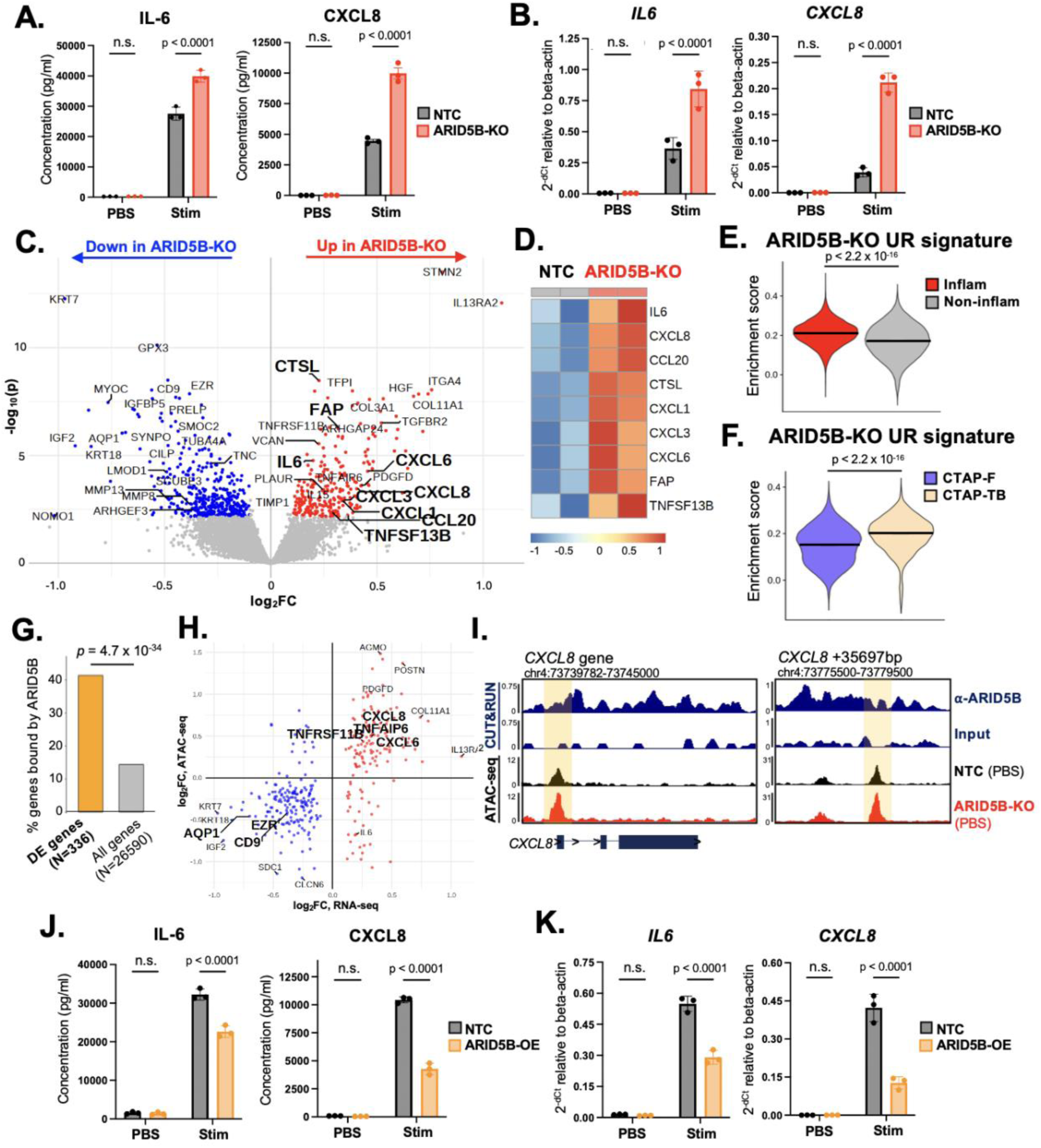
ARID5B suppresses inflammatory fibroblast behavior by reducing inflammatory gene accessibility. **(A)** ELISA for IL-6 and CXCL8 in control (NTC) and ARID5B-deficient (KO) synovial fibroblast supernatants, both unstimulated (PBS) and following stimulation with TNFα, IFNγ, IL-17, and IL1β. Mean ± SD shown. P values from one-way ANOVA with Dunnett multiple comparisons test. **(B)** RT-qPCR for *IL6* and *CXCL8* gene expression in control and ARID5B-deficient synovial fibroblasts following the indicated stimulation conditions. Mean ± SD shown. P values from one-way ANOVA with Dunnett multiple comparisons test. **(C)** Volcano plot showing genes differentially expressed between ARID5B-deficient (KO) and control synovial fibroblasts by bulk RNA-seq. Red points indicate significantly upregulated genes and blue points indicate significantly downregulated genes (*FDR* < 0.05). **(D)** Heatmap of selected inflammatory genes significantly upregulated in ARID5B-deficient (KO) fibroblasts compared to control (NTC). **(E-F)** Violin plots of scores for the gene signature upregulated by ARID5B deletion (ARID5B-KO UR signature) in inflammatory versus noninflammatory synovial fibroblasts **(E)** and in fibroblasts from RA CTAP-F versus CTAP-TB **(F)**. Median expression indicated; p values from two-sided Student’s t-test. **(G)** Enrichment of ARID5B binding at genes differentially expressed upon ARID5B deletion. P value from hypergeometric test. **(H)** Log2-fold change by log2-fold change plot depicting the genes significantly differentially expressed in ARID5B-deficient versus control fibroblasts by RNA-seq (x-axis) and their chromatin accessibility changes in ARID5B-deficient versus control fibroblasts by ATAC-seq (y-axis). Genes upregulated with ARID5B deletion are shown in red; genes downregulated with ARID5B deletion shown in blue (*FDR* < 0.05). **(I)** Track plot of the *CXCL8* gene locus (*left*) and downstream enhancer region (*right*) showing sites of ARID5B binding by CUT&RUN and changes in chromatin accessibility by ATAC-seq between ARID5B-deficient (KO) versus control (NTC) fibroblasts. For CUT&RUN, ARID5B peaks represent averaged signals from all anti-ARID5B antibodies (n=3), with input DNA as negative control. For ATAC-seq, peaks in each condition represent averaged signals from two unstimulated (PBS-treated) synovial fibroblast cell lines (n=2). **(J)** ELISA for IL-6 and CXCL8 in control (NTC) and ARID5B-overexpressing (OE) synovial fibroblast supernatants, both unstimulated (PBS) and following TNFα, IFNγ, IL-17, and IL1β stimulation. Mean ± SD shown. P values from one-way ANOVA with Dunnett multiple comparisons test. **(K)** RT-qPCR for *IL6* and *CXCL8* gene expression in control (NTC) and ARID5B-overexpressing (OE) synovial fibroblasts following the indicated stimulation conditions. Mean ± SD shown. P values from one-way ANOVA with Dunnett multiple comparisons test.

To comprehensively assess the transcriptional effects of ARID5B deletion, we performed bulk RNA-seq on ARID5B-deficient synovial fibroblasts after stimulation with either TNFα alone or in combination with IL-17, IFNγ, and IL1β. We found that in addition to *IL-6* and *CXCL8*, ARID5B loss increased the gene expression of multiple CC/CXC chemokines as well as *TNFRSF13B* (BAFF), all key inflammatory factors implicated in RA (**Figure 3C-D**). We created a signature comprising all genes significantly upregulated in synovial fibroblasts upon ARID5B deletion (*FDR* < 0.05) and assessed its enrichment in the inflammatory and invasive fibroblast subsets defined in the AMP RA/SLE Network Phase 2 synovial tissue dataset (**Figure 1B**). Notably, the gene signature is significantly upregulated among inflammatory fibroblasts compared to noninflammatory fibroblasts (*p* < 2.2 x 10^-16^, **Figure 3E**), suggesting that ARID5B loss causes fibroblasts to retain a more inflammatory transcriptional state. Furthermore, this signature is significantly increased in fibroblasts from CTAP-TB versus fibroblasts from CTAP-F (**Figure 3F**), and it is also enriched in fibroblasts derived from other leukocyte-containing CTAPs, including CTAP-M, CTAP-TM, CTAP-TF, and CTAP-EFM (**Figure S3B**). Thus, the ARID5B-downregulated inflammatory gene signature also delineates differences between RA tissue subtypes.

We next tested whether the genes transcriptionally regulated by ARID5B correspond to genes directly bound by ARID5B. Indeed, among the genes differentially expressed in ARID5B-deficient synovial fibroblasts (*FDR* < 0.05), we observe a significant enrichment for genes identified to be bound by ARID5B from CUT&RUN sequencing (*p* = 4.7 x 10^-34^, **Figure 3G**). These differentially expressed genes are therefore likely to be directly regulated by ARID5B binding. We also compared changes in synovial fibroblast gene expression and chromatin accessibility following ARID5B deletion. Genes differentially expressed in ARID5B-deficient fibroblasts show highly correlated changes in chromatin accessibility (*r* = 0.60, **Figure 3H**), suggesting that chromatin-level gene regulation by ARID5B drives altered expression of ARID5B target genes. Notably, both the *CXCL8* promoter region and a distal downstream enhancer region are bound by ARID5B (**Figure 3I**), with loss of ARID5B leading to an increase in chromatin accessibility at both sites (**Figure 3I**). These regulatory changes explain the increased induction of CXCL8 gene and protein by ARID5B-deficient fibroblasts upon cytokine treatment (**Figure 3A-B**). Collectively, these data suggest that ARID5B binds to key proinflammatory target genes and diminishes their chromatin accessibility, thereby suppressing the synovial fibroblast inflammatory state.

To determine whether ARID5B overexpression conversely blunts the inflammatory activation of synovial fibroblasts, we used CRISPRa to overexpress ARID5B in primary synovial fibroblasts (**Figures S3C and S3D**). Upon combined TNFα, IFNγ, IL-17, and IL1β stimulation, ARID5B-overexpressing synovial fibroblasts express significantly lower levels of IL-6 and CXCL8 relative to their wild-type counterparts, both at the gene and protein level (**Figures 3J and 3K**). ARID5B overexpression also blunts the induction of *IL6* and *CXCL8* in fibroblasts exposed to TNFα alone (**Figure S3E**). Thus, ARID5B is both necessary and sufficient for dampening key molecules that define the fibroblast inflammatory response.

### ARID5B promotes fibroblast invasiveness

Given that ARID5B binds key invasive genes and loss of ARID5B decreases the accessibility of invasive gene loci, we tested whether ARID5B functionally promotes fibroblast invasiveness. We used a transwell invasion assay to evaluate the relative rates at which synovial fibroblasts invade through a Matrigel-coated porous membrane towards a chemoattractant gradient of PDGF-BB^22^. Whereas PDGF-BB significantly increases invasion by wild-type synovial fibroblasts, ARID5B-KO synovial fibroblasts show markedly impaired invasion (**Figures 4A, 4B, and S4A**). We sought to identify the ARID5B-regulated genes responsible for mediating differential fibroblast invasiveness. By bulk RNA-seq, we observed a number of adhesion molecules (*CD9*, *TSPAN2*) and regulators of cell motility and cytoskeleton organization (*EZR*, *AQP1*, *TUBA4A*, *SYNPO*, *KRT*s) to be downregulated in ARID5B-deficient fibroblasts (**Figure 4C**). Interestingly, *TNFRSF11B* (OPG), a key inhibitor of RANKL, was upregulated in ARID5B-deficient fibroblasts, suggesting that ARID5B loss also prevents RANKL-mediated osteoclast activation and bone resorption.

**Figure 4.**
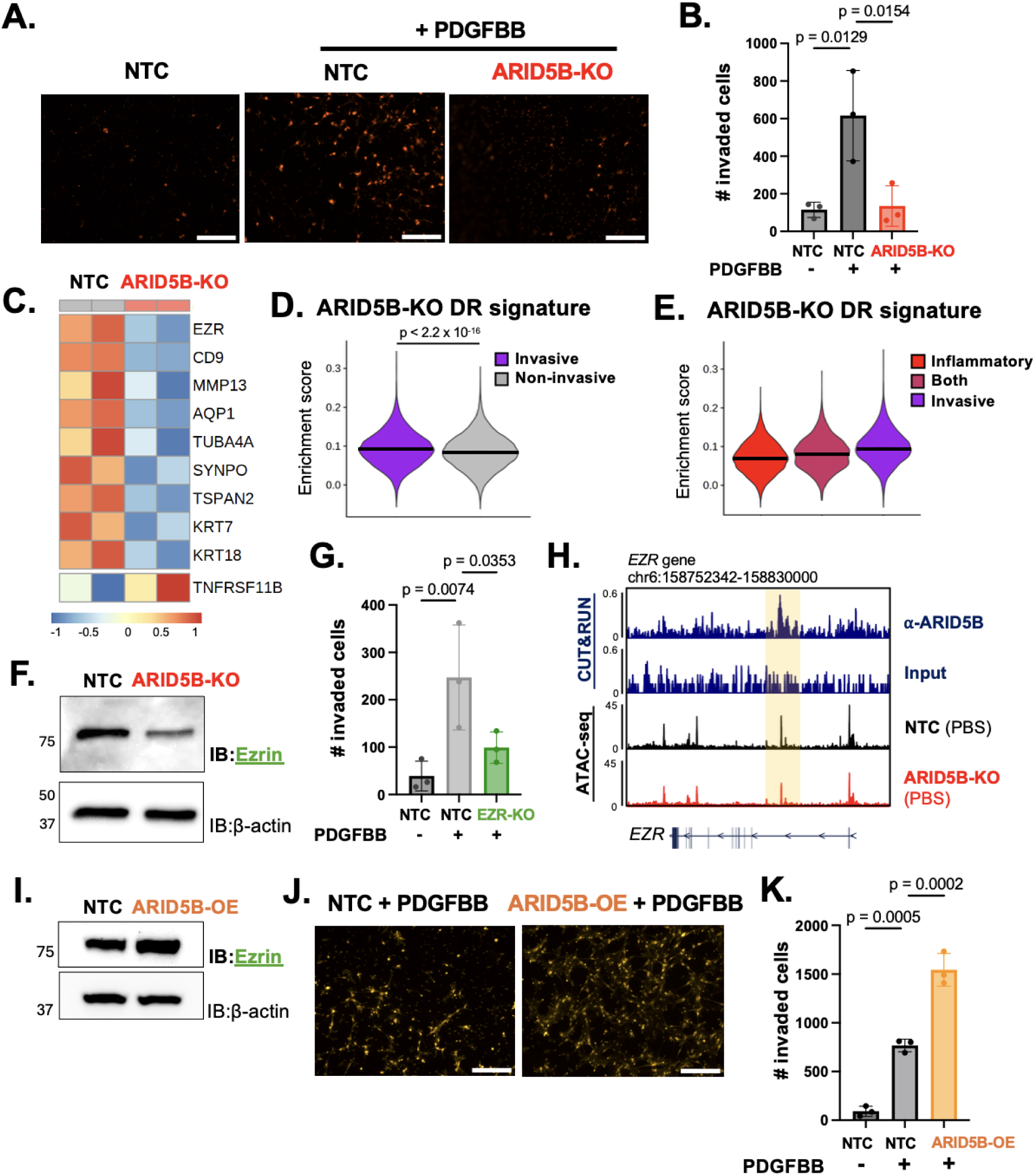
ARID5B promotes fibroblast invasiveness by increasing invasive gene accessibility. **(A)** Representative microscopic fields of invading control and ARID5B-deficient (KO) synovial fibroblasts following PDGF-BB stimulation and propidium iodide staining. Unstimulated fibroblasts shown as negative control. Scale bars represent 250 μm. **(B)** Quantitation of total cellular invasion by control (NTC) and ARID5B-deficient (KO) fibroblasts under the indicated stimulations. Mean ± SD shown. P values from one-way ANOVA with Dunnett multiple comparisons test. **(C)** Heatmap of selected genes regulating cell adhesion, motility, and invasion significantly downregulated in RNA-seq of ARID5B-deficient (KO) versus control (NTC) synovial fibroblasts (*FDR* < 0.05). **(D-E)** Violin plots of scores for the gene signature downregulated by ARID5B deletion (ARID5B-KO DR signature) in invasive versus non-invasive synovial fibroblasts **(D)** and in invasive fibroblasts versus inflammatory and mixed inflammatory-invasive fibroblasts **(E)**. Median expression indicated; p values from two-sided Student’s t-test. **(F)** Immunoblot for Ezrin in control (NTC) and ARID5B-deficient (KO) synovial fibroblast lysates. Beta-actin shown as cell lysate loading control. **(G)** Quantitation of total cellular invasion by control (NTC) and EZR-deficient (KO) synovial fibroblasts under the indicated stimulations. Mean ± SD shown. P values from one-way ANOVA with Dunnett multiple comparisons test. **(H)** Track plot of the *EZR* gene locus showing sites of ARID5B binding by CUT&RUN and differences in chromatin accessibility by ATAC-seq between ARID5B-deficient (KO) versus control (NTC) synovial fibroblasts. For CUT&RUN, ARID5B peaks represent averaged signals from all anti-ARID5B antibodies (n = 3), with input DNA as negative control. For ATAC-seq, peaks in each condition represent averaged signals from two unstimulated (PBS-treated) synovial fibroblast cell lines (n=2). **(I)** Immunoblot for Ezrin in control (NTC) and ARID5B-overexpressing (OE) synovial fibroblasts. Beta-actin shown as cell lysate loading control. **(J)** Representative microscopic fields of invading control (NTC) and ARID5B-overexpressing (OE) synovial fibroblasts following PDGF-BB stimulation and propidium iodide staining. Scale bars represent 250 μm. **(K)** Quantitation of total cellular invasion by control (NTC) and ARID5B-overexpressing (OE) fibroblasts under the indicated stimulations. Mean ± SD shown. P values from one-way ANOVA with Dunnett multiple comparisons test.

To parallel our studies of the genes upregulated by ARID5B deletion (**Figure 3**), we created a gene signature comprising all genes downregulated in ARID5B-deficient fibroblasts and assessed enrichment of this signature in invasive fibroblasts and all other fibroblasts from the AMP RA/SLE Network Phase 2 dataset. At the single-cell level, invasive fibroblasts comprise large proportions of sublining *CD74^hi^* (F-5) and POSTN^+^ (F-3) fibroblasts, as well as a subset of lining *PRG4^+^CLIC5^+^* fibroblasts (F-0) (**Figure S4B**). These invasive fibroblasts show significantly higher enrichment for expression of genes downregulated upon ARID5B deletion, whether relative to all non-invasive fibroblasts (**Figure 4D**) or relative to inflammatory fibroblasts and fibroblasts exhibiting an intermediate combination of inflammatory and invasive states (**Figure 4E**). Thus, loss of ARID5B inhibits the invasive transcriptional state of fibroblasts. In contrast, genes upregulated by ARID5B deletion are enriched both in inflammatory fibroblasts compared to all noninflammatory fibroblasts (**Figure 3E**) and inflammatory fibroblasts relative to invasive fibroblasts and fibroblasts with an intermediate inflammatory-invasive state (**Figure S4C**).

Notably, we found that Ezrin (EZR), an actin cytoskeleton regulator and established driver of cell invasion^50, 51^, is reduced upon ARID5B deletion both at the gene and protein level (**Figures 4C, S4C, and 4F**). While many other genes, including MMPs (MMP13), tetraspanins (CD9), cadherins (CDH11), and EMT regulators (VIM and TWIST1), play roles in controlling fibroblast invasion and are suggested by CUT&RUN or RNA-seq to be regulated by ARID5B, we confirmed that loss of Ezrin alone is sufficient to impair synovial fibroblast invasion (**Figure 4G**). Moreover, upon joint analysis of CUT&RUN and ATAC-seq peaks at the *EZR* locus, we observed that the intronic region of *EZR* is physically bound by ARID5B (**Figure 4H**), with loss of ARID5B decreasing chromatin accessibility at this locus (**Figure 4H**). Thus, ARID5B likely directly regulates Ezrin expression by binding at the *EZR* locus and enhancing *EZR* chromatin accessibility. Indeed, upon ARID5B overexpression, Ezrin gene and protein expression is increased (**Figures S4D and 4I**), and ARID5B-overexpressing synovial fibroblasts invade significantly more efficiently compared to wild-type controls (**Figures 4J and 4K**). Collectively, these findings suggest that ARID5B drives fibroblast invasiveness at least in part by increasing chromatin accessibility of the *EZR* gene locus.

### Arid5b overexpression in fibroblasts drives a shift from inflammatory towards erosive disease

Next, we evaluated how perturbation of ARID5B in fibroblasts affects inflammatory and invasive fibroblast activation *in vivo*. We utilized the K/BxN serum transfer-induced arthritis (STIA) model, in which arthritogenic serum from K/BxN mice is transferred to naïve recipient mice to induce arthritis (**Figure 5A**). As in human RA, murine fibroblasts in STIA play key roles in perpetuating both synovial inflammation and tissue destruction^52, 53^. Whereas inflammatory disease, paw swelling, and fibroblast expression of cytokines such as *Il6* peak during days 7-9 of STIA^40, 54^, bone and cartilage erosion and fibroblast expression of mediators such as *Ezr* peak during days 13-15^55, 56^ (**Figures 5A and S5A**). These observations suggest that fibroblast-driven inflammation precedes and likely triggers fibroblast-mediated tissue damage in STIA, recapitulating the processes observed in RA. Additionally, we confirmed that *Arid5b* is expressed in mouse synovial fibroblasts and is upregulated during peak inflammatory disease in STIA (**Figure 5B**), suggesting it may directly govern pathologic fibroblast functions.

**Figure 5.**
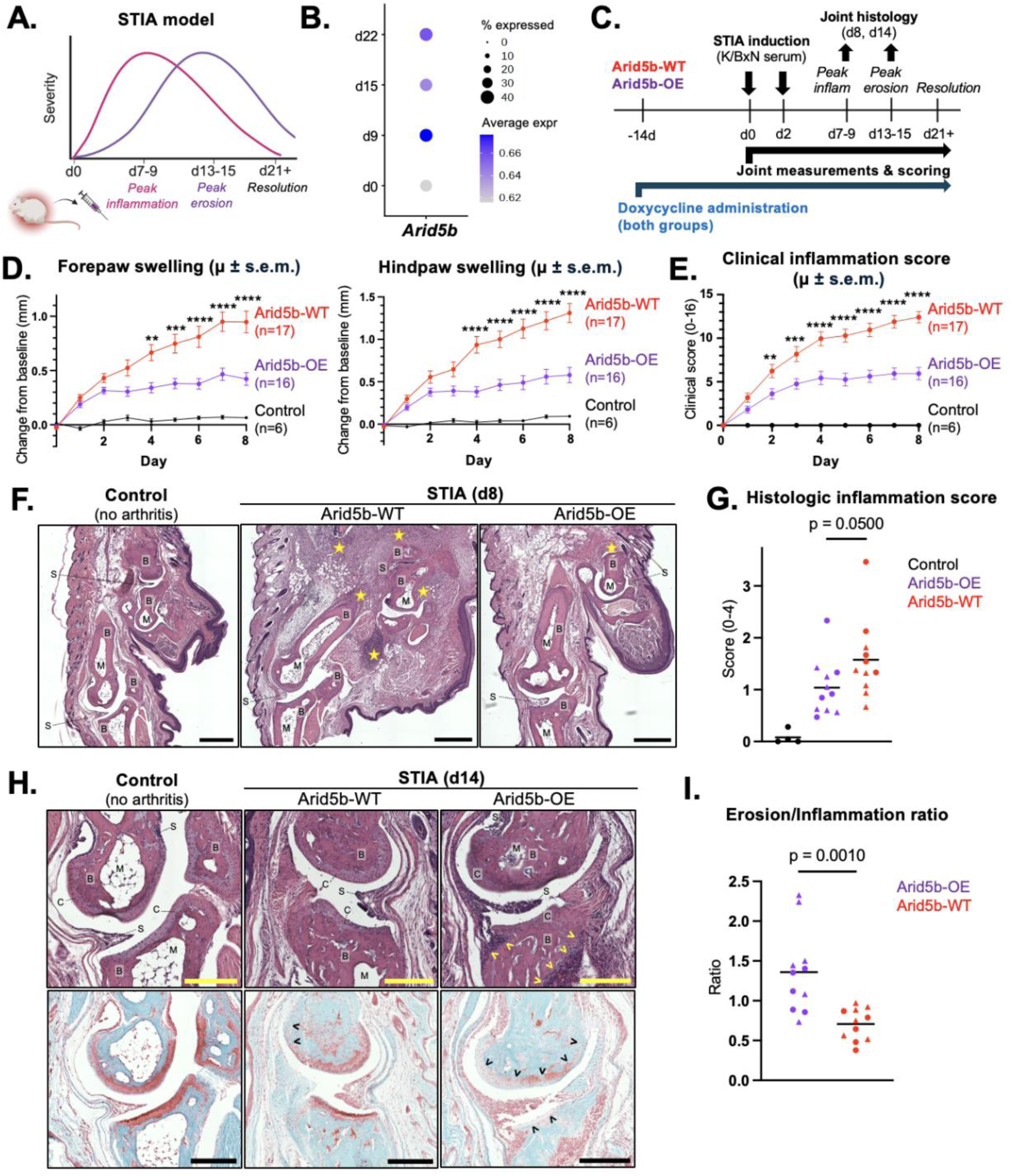
Arid5b-overexpressing fibroblasts shift arthritis pathology from inflammation towards erosion. **(A)** Schematic showing the kinetics of joint swelling and inflammation during K/BxN serum transfer-induced arthritis (STIA). **(B)** Dot plot depicting mean and proportion of *Arid5b* expression in mouse synovial fibroblasts at the indicated STIA timepoints. **(C)** Experimental timeline for the initiation of doxycycline and STIA in Arid5b-OE mice and Arid5b-WT littermates. **(D-E)** Caliper measurements **(D)** and clinical scoring **(E)** of paw swelling and inflammation in Arid5b-OE mice (n = 16) and Arid5b-WT littermates (n = 17). Nonarthritic control mice (n = 6) received no K/BxN serum. Mean ± SEM shown at each timepoint. **p < 0.01; ***p < 0.001; ****p < 0.0001 by two-way repeated measures ANOVA with Šídák’s multiple comparisons test. **(F)** Representative H&E staining of forepaw joints in nonarthritic control mice and arthritic Arid5b-WT and Arid5b-OE mice (STIA day 8). Scale bars represent 500 μm. Asterisks indicate regions of inflammatory infiltrate and arrowheads indicate regions of tissue erosion or damage. B, bone; C, cartilage; M, marrow cavity; S, synovium. **(G)** Histologic inflammation scores for H&E-stained forepaw sections from nonarthritic control mice (n = 4) and arthritic Arid5b-OE and Arid5b-WT mice (circles: n = 5 from STIA day 8; triangles: n = 6 from STIA day 14 per group). P value from one-way ANOVA. **(H)** Representative matched H&E and safranin O (SafO) staining of interphalangeal joints in nonarthritic control, arthritic Arid5b-WT, and arthritic Arid5b-OE mice (STIA day 14). Scale bars represent 250 μm. Yellow arrowheads in H&E images indicate regions of bone erosion; black arrowheads in SafO images indicate areas of cartilage damage (proteoglycan loss or erosion). See (F) for labels. **(I)** Ratio of histologic erosion to inflammation scores for arthritic Arid5b-OE and Arid5b-WT mice (circles: n = 5 from STIA day 8; triangles: n = 6 from STIA day 14 per group). P value from two-sided Student’s t test.

To test the effects of Arid5b on fibroblast behavior during STIA, we generated mice that overexpress *Arid5b* in a fibroblast-specific fashion via a tetracycline-on (Tet-On) system. We crossed Pdgfra^rtTA/+^ mice, which express reverse tetracycline-regulated transactivator (rtTA) under control of the fibroblast-selective *Pdgfra* gene promoter, with Arid5b^tetO/tetO^ mice, which harbor a tetracycline-response element (tetO) upstream of the *Arid5b* start codon^33^ (**Figure S5B**). Doxycycline triggers rtTA binding to the tetO element, enhancing Arid5b gene transcription in fibroblasts. Following two weeks of doxycyline administration, synovial tissue expression of *Arid5b* is significantly increased in Arid5b-OE mice (Pdgfra^rtTA/+^Arid5b^tetO/tetO^) relative to Arid5b-Het (Pdgfra^rtTA/+^Arid5b^tetO/+^) and Arid5b-WT (Pdgfra^+/+^) littermates (**Figure S5C**). We subsequently initiated STIA and performed joint measurements, scoring, and histology throughout the disease course (**Figure 5C**).

Notably, the peak severity of joint inflammation is significantly blunted in Arid5b-OE mice compared to Arid5b-WT mice, both by caliper measurement and clinical assessment of paw swelling (**Figures 5D and 5E**). Correspondingly, histologic evaluation of paw joints during the peak of inflammatory disease (day 8) confirmed decreased synovitis and inflammatory cell infiltrates in Arid5b-OE relative to Arid5b-WT mice (**Figure 5F**). Arid5b-OE mice also maintain reduced paw swelling over the subsequent disease course (days 9-25) (**Figures 5G, S5D, and S5E**). Together, these findings indicate that Arid5b restrains the inflammatory activation of fibroblasts and in turn, their ability to drive continued synovial inflammation in arthritic mice.

We next assessed how Arid5b overexpression affected erosive disease. In patients with RA, the severity of bone and cartilage damage is prominently driven by the severity of inflammatory disease^24, 25^. Similarly, in Arid5b-WT mice, histologic erosion scores increase linearly with histologic inflammation scores (**Figure S5F**). Given that the Arid5b-OE mice had reduced inflammatory disease burden compared to Arid5b-WT mice (**Figures, 5D, 5E, 5F, and 5G**), we predicted they would likewise show diminished erosive disease compared to Arid5b-WT mice. However, despite decreased inflammation, Arid5b-OE mice exhibited bone and cartilage damage of a comparable degree to their Arid5b-WT counterparts over the arthritis course (days 8 and 14) (**Figure S5G**). Histologic evaluation of paw joints at the peak of erosive disease (day 14) revealed extensive loss of proteoglycan in articular cartilage as well as synovial pannus formation and infiltration of subchondral bone in both Arid5b-WT and Arid5b-OE mice (**Figure 5H**). When normalized for differences in joint inflammation, Arid5b-OE mice exhibited a significant shift towards erosive disease from inflammatory disease relative to Arid5b-WT mice (**Figure 5I**). Altogether, our work points to ARID5B as an axis that enables an activated inflammatory fibroblast to acquire invasive capabilities, which in turn dictates the inflammatory-to-degradative shift in fibroblast-driven tissue pathology that defines RA (**Figure S5H**).

## Discussion

Here, we show that ARID5B is a key transcription factor highly expressed in inflammatory fibroblasts in RA and upregulated by inflammatory cytokines implicated in RA pathology. In cytokine-activated fibroblasts, ARID5B complexes with histone editors and binds to both inflammatory and invasive gene loci. By decreasing the chromatin accessibility of pro-inflammatory genes while increasing the chromatin accessibility of motility and invasion-promoting genes, ARID5B limits the inflammatory activity of fibroblasts while driving their invasiveness. In a mouse model of inflammatory arthritis, fibroblast-specific Arid5b overexpression increases the relative severity of erosive disease over inflammatory disease.

Understanding of the mechanisms by which pathologic fibroblast functions are regulated is still in its early stages. As potent regulators of cell state and function, TFs have provided increasing insight into the gene expression programs driving fibroblast pathology. Thus far, however, these TFs have largely been observed to regulate a singular fibroblast function or phenotype. Notably, PU.1 (*SPI1*), upregulated in fibrotic diseases, drives myofibroblast differentiation and fibrogenesis^30^, whereas ETS1 promotes a tissue degradative fibroblast phenotype with induction of RANKL and MMP expression^31^. Likewise, effectors of cytokine signaling, such as STAT4^18^, and transducers of Notch-induced fibroblast differentiation, such as RBPJ^40^, predominantly drive the inflammatory activation of fibroblasts.

In contrast, we observe that ARID5B regulates the inflammatory and invasive behaviors of fibroblasts in tandem, governing a transition from fibroblast-driven tissue inflammation to fibroblast-mediated bone and cartilage destruction. Our work suggests that ARID5B, when upregulated by inflammatory stimuli, acts as a feedback inhibitor which epigenetically suppresses the ability of the fibroblast to undergo further inflammatory activation and in turn shifts the fibroblast towards a more invasive state. This axis of ARID5B upregulation and function could be akin to how checkpoint molecules such as PD-1 and CTLA4 are transiently induced upon leukocyte stimulation to contain the primary activation response^57, 58^. Additionally, it provides insight into how the highly pro-inflammatory, cytokine-rich tissue microenvironment observed in RA subsequently gives rise to a robust tissue degradative program that erodes cartilage and bone.

Unlike previously described TFs in fibroblasts, which directly activate or repress target gene transcription, we find that ARID5B instead regulates a broad range of inflammatory and invasive gene loci through reciprocal control at the epigenetic level. In RA fibroblasts, ARID5B complexes with histone editors and simultaneously dampens the chromatin accessibility of inflammatory genes (such as *CXCL8* and other CC/CXC chemokines) while enhancing the accessibility of invasive genes (such as *EZR* and other cell adhesion and cytoskeleton regulators).

Altered epigenetic regulation of synovial fibroblasts is known to contribute prominently to RA pathology. Synovial fibroblasts isolated from RA joints retain inflammatory and invasive characteristics when cultured *ex vivo* and upon adoptive transfer into naive mice^54, 59^. Moreover, epigenetic processes have been hypothesized to reinforce the maintenance of inflammatory and invasive gene expression programs in pathologic fibroblasts. For instance, synovial fibroblasts from RA patients have been shown to exhibit increased levels of activating histone marks and decreased levels of repressive histone marks at *IL6*, *CXCL8*, and *MMP* gene loci^60, 61^. Single-cell studies have further implicated the chromatin accessibility levels of several inflammatory genes, including *IL-6*, *CXCL12*, and *HLA-DR*, and matrix degradative and invasive genes, such as MMPs, as prominent features of pathologic fibroblast populations in RA^62^. By inducing reciprocal changes in chromatin accessibility at inflammatory and invasive gene loci, ARID5B mechanistically explains how epigenetic regulation shapes both the inflammatory and invasive potential of an activated fibroblast.

Of note, while our work describes a pathway governing an inflammatory-to-invasive shift in pathologic fibroblast state, previous work in murine arthritis models has suggested instead that distinct inflammatory and invasive fibroblast populations arise within separate anatomic locations in arthritis synovium to mediate these functions independently^54^. Specifically, fibroblasts from the synovial sublining layer, expressing fibroblast activation protein alpha (FAPα) and THY1 (CD90), exacerbated the inflammatory severity of arthritis but did not affect damage to cartilage or bone, while lining-layer FAPα^+^THY1^-^ fibroblasts selectively aggravated bone and cartilage damage without impacting inflammation^54^. However, studies of human synovium have demonstrated that RA is associated with predominant expansion of the sublining fibroblast populations, and that in contrast, a predominant expansion of lining fibroblasts is associated with osteoarthritis^17^. Furthermore, among RA patients of varying synovial tissue subtypes, sublining fibroblasts were observed to be enriched in tissues containing higher proportions of inflammatory leukocytes, whereas lining fibroblasts were enriched in leukocyte-poor subtypes^9^. These findings suggest that in chronic inflammation in humans, the anatomically separate sublining and lining synovial fibroblasts may become preferentially activated in distinct disease contexts with inflammatory sublining fibroblasts capable of demonstrating significant invasive signatures.

Previously, ARID5B was largely studied in the context of leukocyte biology. Notably, ARID5B was one of the first risk genes linked to acute lymphoblastic leukemia (ALL)^63, 64^, and it has been shown to regulate B and NK cell development, metabolism, and effector function^33, 48^. Among stromal cell types, ARID5B is comparatively less well-understood, but it has been shown to regulate adipogenesis and chondrocyte differentiation^32, 65^. Additionally, a previous study of TFs in synovial fibroblasts identified ARID5B as a negative regulator of IL-6 production *in vitro*^66^. In addition to IL-6, our work illustrates the ability of ARID5B to regulate a broad range of both inflammatory and invasive effector genes in synovial fibroblasts, including CXCL8, CC/CXC chemokines, Ezrin, MMP13, and other cell adhesion and motility molecules. We further show at the epigenetic, transcriptional, and functional levels that ARID5B broadly controls a shift in the fibroblast state from inflammatory to invasive.

We also find that fibroblast-specific Arid5b overexpression in mice is sufficient to blunt inflammatory disease during arthritis while promoting a relative increase in bone and cartilage erosion. This functional axis may contribute to driving the differences in synovial tissue pathology observed among patients with RA^5, 17, 67^. Indeed, we observe that whereas ARID5B is elevated in synovial tissues rich in activated proinflammatory leukocytes, it is downregulated in tissues harboring a lower proportion of leukocytes. Furthermore, the *ARID5B* locus itself contains SNPs linked to RA risk that overlap active *cis*-regulatory elements which are predicted to modulate *ARID5B* expression. Thus, regulation of ARID5B levels in fibroblasts, and in turn, the propensity of activated fibroblasts to maintain inflammatory responses or acquire invasive characteristics upon stimulation, may directly impact RA pathology and disease progression and underpin the observed heterogeneity in disease phenotypes and clinical presentation. Altogether, our work describes an inflammatory-to-invasive axis driven by ARID5B in fibroblasts and underscores its influence on end-organ pathology. More broadly, our work suggests that any therapeutically successful fibroblast-targeting approach must account for the distinct axes regulating inflammatory and invasive fibroblast behavior.

## Resource availability

### Lead contact

Additional information or requests for resources will be fulfilled by the lead contact, Michael Brenner.

### Materials availability

This study did not generate new unique reagents.

### Data and code availability

The following datasets have been deposited at Zenodo and will be made publicly available upon publication: CUT&RUN sequencing data (https://zenodo.org/records/15312813?preview=1&token=eyJhbGciOiJIUzUxMiJ9.eyJpZCI6Ijcx MWU3ODkwLTJjYmEtNDUzOS1hZmQxLTQyZjhkYTBlNDBhMCIsImRhdGEiOnt9LCJyYW5kb2 0iOiJlNjVlMDAwN2QxN2Q2MWM4ZjM2NTc0ODEzNjkwZGZlYiJ9.k5ktzONPpflzAo52Er44GRH 1LXk5mMXyxzdA5XWYLCS5x8X6hjVjdiPnTQHEXgQA550-JyGEbqtEKitJKi_bqg), RNA sequencing data (https://zenodo.org/records/15313192?preview=1&token=eyJhbGciOiJIUzUxMiJ9.eyJpZCI6IjBj MmQxZWEyLTk3MTEtNDJkMy1iMWNlLTNjYzRjZDA1NTdmMiIsImRhdGEiOnt9LCJyYW5kb20i OiI3YzU5OGVkYmZjMzY4ZDA4YzI2YWNiMmEyOTQzZmY5OSJ9.2Us_8BeGsvjW9x9-1MiE5hI9EJgebhB02CeDty65t8KEnZ8UNYhAzVfszayys0HwA8IXCas2OXdzzJ8CiKkFLw), and ATAC sequencing data (https://zenodo.org/records/15313206?preview=1&token=eyJhbGciOiJIUzUxMiJ9.eyJpZCI6IjI1 NDlkNWVkLWYyMWItNDEzMC05NzA1LTU2ODk0OGVkMGIzMCIsImRhdGEiOnt9LCJyYW5kb20iOiJhMjRjN2MzNmY1ZDRiNTdhNDUzNjcyZGY5YjdkYzg2NCJ9.G9EkPpr4-g86jjbRdH6hO24DJWn7F15r7VFlUEQwGzLXic8bQ9JmyZ3zErm71f9vFl0NvaGkfaXslICDjlzlPg; https://zenodo.org/records/15610388?preview=1&token=eyJhbGciOiJIUzUxMiJ9.eyJpZCI6Ijk3Y jMyYjg5LTFmODgtNGY5ZS1hODQ5LTc1MzYzNTUzZDg1NCIsImRhdGEiOnt9LCJyYW5kb20i OiI1MjQ2ZDBjNGI0Mzc2ZWE5OTg3NzJlMzJjODg4NDQ3YiJ9.rKEcsXEiyXxIMLqA_0YT9lg2M7YlxMka_7Ofe-VqV46wOsVQKCh7UizbTy72u4n0OweT7LBvc9RRdXM1dkrg4w; https://zenodo.org/records/15660468?preview=1&token=eyJhbGciOiJIUzUxMiJ9.eyJpZCI6ImM 0Y2FkMGE2LTQxY2MtNGM2MC1hZjNmLTM3Y2QxNmM1MjQxOSIsImRhdGEiOnt9LCJyYW5kb20iOiJkNmU1YTgzNmZjYjEzMmM1OGM0ZDYzOTc2ZGU5ZmVjMSJ9.WCeFex8Y90HZsaW VyKo4rLchNMRtQoxg5bqDhV_BioUER8lmb75WApUaU5-O7ASAuL2R-Y40omkF0h3bkHtOGQ; https://zenodo.org/records/17316817?preview=1&token=eyJhbGciOiJIUzUxMiJ9.eyJpZCI6IjMx NmViNDQwLWZhZGMtNGNmYi1iNzNiLWQ1YWRhNzBkOWYzZiIsImRhdGEiOnt9LCJyYW5kb 20iOiI3NzJmOTJlMTcyOTJmMDM4Nzc4NzdiNWNhYTRhOGI2YiJ9.SHwvuiFacb7BAxEupaG9r W_Ni3IfVXxPO0vY-3T6Ko9_Q3EIE3_uFkJinHMJ-IAvD0ptVU9Ql0gci7W6cbkDJw)

## Acknowledgements

The authors thank Drs. Jun Yang and Megan Walker for providing Arid5b^tetO^ mice and Dr. Jacqueline Perrigoue for helpful discussions. This work is supported by NIH grants F30AI174699, T32GM007753, and T32GM144273 (to A.E.Z.); K08AR083513, T32AR007530, and P30AR070253 (to A.A.M.); and R01AR0637039, P01AI148102 (to M.B.B.). A.A.M. also received grant support from the Rheumatology Research Foundation.

## Author contributions

A.E.Z. and M.B.B. conceived the study. A.E.Z., S.K., G.F.M.W., and M.L.F. performed the experiments and analyzed the data. A.E.Z., M.B.B., and A.A.M. drafted the manuscript with input from all authors. M.B.B. and A.A.M. supervised the study.

## Declaration of interests

M.B.B. is on the scientific advisory board of AbbVie and Moderna, a consultant to 4F0 Ventures and a founder of Mestag Therapeutics.

## Methods

### Cell culture

Primary synovial fibroblast cell lines were derived from the synovial tissues of adult RA patients (male and female) diagnosed per 2010 ACR/EULAR classification criteria, following biopsy, synovectomy, or arthroplasty. Cells were cultured in a humidified 10% CO2 incubator at 37°C with Dulbecco’s Modified Eagle Medium (DMEM) supplemented with 10% fetal bovine serum (FBS) (Gemini Bio), 1% MEM non-essential amino acids (Gibco), 2% MEM amino acids (Gibco), 100 U/ml penicillin (Gibco), 100 μg/ml streptomycin sulfate (Gibco), 2mM L-Glutamine (Gibco), 55uM 2-Mercaptoethanol (Gibco), 10ug/mL Gentamicin (Gibco), and 10 mM HEPES (Gibco). Fibroblasts were maintained for up to 10 passages for experiments.

### Animals

Animals were housed in AAALAC-accredited facilities at Brigham and Women’s Hospital (BWH). All studies were performed with approval by the BWH Institutional Animal Care and Use Committee (IACUC). Mice were maintained in pathogen-free, humidity-controlled, and temperature-controlled housing with 12-hour day/night cycles and regular veterinary monitoring. Arid5b^tetO^ mice were generated by fertilizing C57BL/6 ova with cryopreserved Arid5b^tetO/tetO^ sperm (gift from Dr. Jun Yang, St. Jude Children’s Research Hospital, Memphis, TN^33^). These mice were crossed with Pdgfra^rtTA/+^ mice (Jackson Labs; strain #034459) to generate Pdgfra^rtTA/+^Arid5b^tetO/tetO^ mice (Arid5b-OE mice) and Pdgfra^+/+^ littermate controls (Arid5b-WT mice). Genotyping was performed using TransnetYX. Arid5b overexpression was induced by administering doxycycline chow (625 mg/kg; Bio-Serv) for two weeks prior to the beginning of each experiment. Overexpression of Arid5b in Arid5b-OE mice was confirmed by RT-qPCR of dissected knee synovial tissue. Both Arid5b-OE and Arid5b-WT mice remained on doxycycline chow for the duration of each arthritis experiment.

### Coimmunoprecipitation

Primary human synovial fibroblasts were washed in cold PBS, followed by addition of IP lysis buffer [50 mM Tris pH 8, 2.5% glycerol, 150 mM NaCl, 0.5% Triton-X 100, 1mM sodium orthovanadate, 1mM phenylmethylsulfonyl fluoride, 1mM beta-glycerophosphate, and 1mM sodium pyrophosphate, supplemented with cOmplete protease inhibitor cocktail (Roche)].

Samples were lysed for 30min at 4°C followed by centrifugation at 15,000 rpm for 15min at 4°C. Immunoprecipitation was performed overnight at 4°C using Protein A resins (Invitrogen Dynabeads) along with an anti-ARID5B antibody (Novus Biologicals, NBP1-83622) or rabbit IgG isotype control (Cell Signaling Technologies, 2729).

### Immunoblot

Cells were lysed for 1h at 4°C in RIPA buffer supplemented with cOmplete protease inhibitor cocktail (Roche). Cell lysate concentrations were quantified using a BCA protein assay kit (Thermo Fisher). Sample proteins were separated by SDS-PAGE on Mini-PROTEAN TGX gels (anyKD or 7.5%, Bio-Rad) and transferred using the Trans-Blot Turbo Transfer System (Bio-Rad) onto 0.2µM PVDF membranes. Membranes were blocked for 1h at room temperature with 5% nonfat dry milk in TBST (0.1% Tween 20) and incubated overnight with primary antibodies diluted in blocking buffer at 4°C. Membranes were subsequently incubated with HRP-conjugated secondary antibodies for 1h at room temperature, developed using the Clarity Western ECL Substrate (Bio-Rad), and imaged on a ChemiDoc Touch MP System (Bio-Rad). Primary antibodies used included: anti-ARID5B #1 (Novus Biologicals, NBP1-83622), anti-ARID5B #2 (Novus Biologicals, NBP3-13006), anti-HDAC1 (Cell Signaling Technologies, 34589), anti-HDAC2 (Cell Signaling Technologies, 57156), anti-Ezrin (Cell Signaling Technologies, 3145), and anti-beta-actin (Millipore Sigma, A5441). Secondary antibodies used included: HRP-conjugated goat anti-mouse IgG (Jackson ImmunoResearch, 115-035-146), HRP-conjugated donkey anti-rabbit IgG (Jackson ImmunoResearch, 711-035-152).

### CRISPR/Cas9 gene deletion

sgRNA sequences against each target gene were designed using the CHOPCHOP algorithm^68^. (NTC sequences: GGAGAGGGCCCGCGAACUCA, CUGACGUGUCUGAAAUGAGU, GACGCCUUGCCCGGCUCACA; ARID5B sequences: UGUAGGCUGAUCAUCCGUAG, CACUCGCUUGUGAUGCAAAG, UAAGGUCCGUGCAAGCCACA, GUUUCAGCAUCGAGCGGUAC, CGAGCGGUACCGGCAGUACU; EZR sequences: UGUGGCAUGCGGAACACCGU, AUGAGUUGUAUAUGCGCCGC, UAGCUCACCGGCUCGUACAC). To form Cas9/sgRNA ribonucleoprotein (RNP) complexes, 20uM recombinant SpCas9 2NLS Nuclease (Synthego) was combined with 80uM sgRNA (Synthego) and incubated at room temperature for 15min. To maximize deletion efficiency, we pooled 2-3 sgRNA sequences against each target except where indicated in individual figure panels. For RNP nucleofection, primary human synovial fibroblasts were resuspended in P2 Primary Cell Nucleofector Solution (Lonza; 17µl per reaction) and combined with the assembled Cas9/sgRNA RNPs (3µl per reaction) in a nucleocuvette vessel plate (Lonza). Cells underwent nucleofection using the EN-150 program on a 4D-Nucleofector X Unit (Lonza) and were reseeded and allowed to recover and expand for at least seven days prior to experimental studies. Gene deletion was confirmed by immunoblot.

### CRISPR activation (CRISPRa)

Primary human synovial fibroblasts that stably express the fusion proteins dead Cas9 (dCas9)-VP64 (Addgene) and MS2-p65-HSF1 (Addgene) were transduced with lentiviruses encoding CRISPRa sgRNAs for each target gene in the pXPR_502 backbone (Addgene). We used the following sgRNA sequences from the Calabrese P65-HSF CRISPRa library (*NTC*: AAGAATTAGGCACGGTTACT; *ARID5B*: GAGTTGCCAGACTGGCAGGG). Transductions were performed at an MOI of 0.3 in the presence of 10µg/ml polybrene, with centrifugation at 930g for 2h at 30°C. Transduced fibroblasts were allowed to recover and expand for four days and subsequently underwent dual blasticidin and puromycin selection for at least 14 days prior to experimental studies. Gene overexpression was confirmed by RT-qPCR and immunoblot.

### RT-qPCR

For cytokine stimulation experiments, primary human synovial fibroblasts were stimulated with either TNFα (10 ng/ml, 16h), TNFα+IFNγ+IL-17+IL1β (TNFα: 0.1 ng/ml, IFNγ: 1 ng/ml, IL-17: 1 ng/ml, IL1β: 1 pg/ml, for 16h), or an equivalent volume of PBS (unstimulated). Total RNA from human fibroblasts or mouse synovial tissue was extracted using the RNeasy Mini Kit (Qiagen). Reverse transcription and cDNA synthesis were performed with QuantiTect RT Kit (Qiagen) using a BioRad T100 Thermal Cycler. RT-qPCR was conducted using Brilliant III Ultra-Fast SYBR Green QPCR Master Mix (Agilent Technologies) with the following primers: *ACTB* (F: CACCATTGGCAATGAGCGGTTC, R: AGGTCTTTGCGGATGTCCACGT), *ARID5B* (F: GAATTAGGCGGTAATCCTGGGAG, R: TCCGAGGTTTGATTGGAGGCAG), *IL6* (F: CCACTCACCTCTTCAGAACG, R: CATCTTTGGAAGGTTCAGGTTG), *CXCL8* (F: ACTGAGAGTGATTGAGAGTGGAC, R: AACCCTCTGCACCCAGTTTTC), *EZR* (F: ATCGAGGTGCAGCAGATGAAGG, R: CGCAGCATCAACTCCTCCTTCT), mouse *Actb* (F: CATTGCTGACAGGATGCAGAAGG, R: TGCTGGAAGGTGGACAGTGAGG), and mouse *Arid5b* (F: AATCTTGTCCCTTGGCGACT, R: CAGAATGCGCCCATTTGACA). Gene expression was quantitated using the ΔΔCT method, with *ACTB* or *Actb* mRNA as the internal control.

### ELISA

For cytokine stimulation experiments, primary human synovial fibroblasts were stimulated with either TNFα (10 ng/ml, 16h), TNFα+IFNγ+IL-17+IL1β (TNFα: 0.1 ng/ml, IFNγ: 1 ng/ml, IL-17: 1 ng/ml, IL1β: 1 pg/ml, for 16h), or an equivalent volume of PBS (unstimulated). Culture supernatants were harvested from fibroblasts for assessment of IL-6 and CXCL8 concentrations by DuoSet ELISA (R&D; IL-6: DY206; CXCL8: DY208), per manufacturer instructions. Briefly, 96-well plates were coated with a capture antibody and blocked to prevent nonspecific binding. Sample supernatants and standards were added, followed by incubation with detection antibody and a streptavidin-HRP conjugate. TMB substrate solution was added for color development and stopped with a 2N H_2_SO_4_ acid solution. Sample and standard absorbances were measured on a Spectra Max microplate reader with mean absorbance of the zero standard subtracted from each reading. Standard curves were generated from the absorbances of all standards using a four-parameter logistic (4-PL) curve-fit (GraphPad Prism). Sample concentrations were interpolated from the standard curve.

### Transwell invasion assay

We adapted the transwell invasion assay described Kiener et al.^22^ as follows. Matrigel invasion chambers (BioCoat GFR, Corning) were rehydrated and invasion medium (DMEM, supplemented with 0.1% serum albumin, 2 mM L-glutamine, 10 μM non-essential amino acids, 100 U/ml penicillin, and 100 μg/ml streptomycin sulfate) containing PDGF-BB (StemCell Technologies) as a chemoattractant was added to the bottom wells of the chambers. Based on the donor primary synovial fibroblast cell line and passage, we used 100-300 ng/ml PDGF-BB in order to elicit at least a five-fold increase in fibroblast invasion above untreated control synovial fibroblast cell lines. 2.5 × 10^4^ synovial fibroblasts were plated in the top chamber and incubated for 16h at 37°C. Then, membranes were fixed and permeabilized with ice-cold methanol. Cells that failed to penetrate the membrane were removed from the top of the membrane using cotton swabs, and cells that invaded to the underside of the membrane were stained with propidium iodide (50 μg/ml propidium iodide in 1 mg/ml D-glucose/PBS). Membranes were imaged by fluorescence microscopy. Nine microscopic fields (10X) per membrane were counted, with each condition repeated in triplicate. Data were expressed as the average number of invaded cells per nine fields.

### RNA-sequencing

#### Sample preparation

Primary human synovial fibroblasts from two RA patients underwent CRISPR deletion with NTC (control) or ARID5B-targeting sgRNAs and were stimulated with the following cytokines: in Experiment 1, TNFα (10 ng/ml, for 4h or 24h), and in Experiment 2, TNFα+IFNγ+IL-17+IL1β (TNFα: 0.1 ng/ml, IFNγ: 1 ng/ml, IL-17: 1 ng/ml, IL1β: 1 pg/ml, for 16h). Cells were lysed in TCL buffer (Qiagen). Sequencing libraries were prepared using the Illumina Smart-Seq2 protocol and sequenced on an Illumina NextSeq 500 instrument using 2x38bp paired-end reads by Broad Clinical Labs.

#### Data analysis

We used kallisto (v0.46.2)^69^ to pseudoalign and quantify reads against the GRCh38 v104 transcriptome. Differential gene expression analyses were performed with limma (v3.50.3)^70^. We fit the following linear model to each gene: *GeneExpression* ∼ *Genotype* + *Cell.line* + *Dataset* + *Time* + *Cell.line:Stim* + *Cell.line:Time*.

*Genotype* refers to whether fibroblasts are wild-type or ARID5B-deficient; *Cell.line* represents two indicator variables, one for each synovial fibroblast cell line used; *Dataset* refers to whether fibroblasts were derived from Experiment 1 or 2; and *Time* represents the duration of fibroblast stimulation. *Cell.line:Stim* and *Cell.line:Time* represent interactions between synovial fibroblast cell line and stimulation type or time, respectively. All genes with *FDR* < 0.05 were considered to be significantly differentially expressed in ARID5B-deficient fibroblasts.

### CUT&RUN sequencing

#### Sample preparation

Primary synovial fibroblasts from an RA patient were stimulated with TNFα+IFNγ+IL-17+IL1β (TNFα: 0.1 ng/ml, IFNγ: 1 ng/ml, IL-17: 1 ng/ml, IL1β: 1 pg/ml) for 16h. CUT&RUN reactions were performed with the CUT&RUN Assay Kit (Cell Signaling Technologies) according to manufacturer instructions. Briefly, 1 x 10^5^ fibroblasts per reaction were collected without fixation, bound to concanavalin A beads, permeabilized with digitonin, and incubated with primary antibodies against the targets of interest overnight at 4°C. The following primary antibodies were used: anti-ARID5B antibody #1 (Novus Biologicals, NBP1-83622), anti-ARID5B antibody #2 (Novus Biologicals, NBP3-13006), anti-ARID5B antibody #3 (Millipore Sigma, HPA015037), anti-Tri-Methyl-Histone H3 (Lys4) (C42D8) positive control (Cell Signaling Technologies, 9751), and anti-rabbit IgG isotype control (CUT&RUN, DA1E) (Cell Signaling Technologies, 66362). Next, protein A/G-micrococcal nuclease (MNase) conjugates were added and the MNase enzyme was activated by calcium to enable selective digestion of DNA complexed to the antibody-bound protein target.

For the input sample, collected fibroblasts were incubated with DNA extraction buffer + Proteinase K + RNase A (Cell Signaling Technologies) for 1h at 55°C with shaking per manufacturer instructions. Cell lysis and chromatin fragmentation were performed by sonication using a VirTis Virsonic 100 Ultrasonic Homogenizer at setting 6 (20% amplitude) with a ⅛ inch probe. We performed 5 sets of 15s pulses and 30s rest on ice in between pulses.

Digested sample and input DNA were purified using the DNA Purification Buffers and Spin Columns kit (Cell Signaling Technologies) according to manufacturer instructions. DNA libraries were prepared and sequenced by the Molecular Biology Core Facilities at Dana-Farber Cancer Institute using 2x150bp paired-end reads to a depth of 10M reads.

#### Data analysis

Reads were aligned to the GRCh38 reference genome (no-alt analysis set) using bowtie2 (v2.5.1)^71^, with --local and --dovetail settings enabled. Read alignments with MAPQ < 10 were excluded. Peak calling was performed with MACS2 (v2.2.7.1)^72^ against both input DNA control and IgG isotype control (q-value < 0.05). Sample quality was assessed using ChIPQC (v1.30.0)^73^. Functional and gene annotation of peaks was performed with ChIPseeker (v1.30.3)^74^. To identify a consensus set of genes bound by ARID5B, we determined the intersection of genes bound using each anti-ARID5B antibody based on peaks called against the input control. We performed Gene Ontology term enrichment analysis on the consensus ARID5B-bound genes using the PANTHER Overrepresentation Test^46^. We used trackplot^75^ to visualize CUT&RUN peaks.

### ATAC-sequencing

#### Sample preparation

Primary synovial fibroblasts from two RA patients underwent CRISPR deletion with NTC (control) or ARID5B-targeting sgRNAs, followed by no treatment (PBS) or by stimulation with TNFα+IFNγ+IL-17+IL1β (TNFα: 0.1 ng/ml, IFNγ: 1 ng/ml, IL-17: 1 ng/ml, IL1β: 1 pg/ml) for 16h.

Libraries were prepared with the ATAC-Seq Kit (Active Motif) using unique dual indices (UDI). Briefly, 1 x 10^5^ fibroblasts per reaction were collected and 1.0 × 104 cryopreserved Drosophila cell nuclei (Active Motif) were added as spike-in controls to each sample. Samples subsequently underwent tagmentation, DNA purification, amplification of tagmented DNA, and SPRI bead cleanup of amplified DNA. Libraries were sequenced by the Molecular Biology Core Facilities at Dana-Farber Cancer Institute using 2x150bp paired-end reads.

#### Data analysis

Sequencing reads were trimmed with TrimGalore (v0.5.0) and aligned to the GRCh38 reference genome (no-alt analysis set) using bowtie2 (v2.5.1)^71^, with --local and --dovetail settings enabled. Read alignments to mitochondrial DNA or with MAPQ < 10 were excluded. For spike-in normalization, reads were aligned to the Drosophila BDGP6 reference genome using bowtie2. Based on the number of uniquely aligning Drosophila reads detected in each sample, normalization factors were calculated and samples were downsized to normalize human read counts. PCR duplicates were removed with Picard MarkDuplicates (v2.24.1). Tn5 shifting was performed with deepTools alignmentSieve (v3.5.2)^76^. Peaks were called with MACS2 (v2.2.7.1)^72^ after merging all ATAC-seq samples. Differential accessibility analyses were performed with csaw (v1.28.0)^77^. Sample reads were counted in each peak with low abundance peaks (logCPM < -3) excluded. Samples were normalized using a nonlinear loess-based approach (Method IV as described^78^). We fit the following model to each genomic region tested: *Accessibility* ∼ *Genotype* + *Cell.line* + *Stim*.

*Genotype* refers to whether fibroblasts are wild-type or ARID5B-deficient; *Cell.line* represents two indicator variables, one for each synovial fibroblast cell line used; *Stim* refers to whether fibroblasts are untreated or stimulated with TNFα+IFNγ+IL-17+IL1β. All peaks with FDR < 0.15 were considered to be significantly differentially accessible in ARID5B-deficient fibroblasts.

Functional and gene annotation of peaks was performed with ChIPseeker (v1.30.3)^74^. Peaks were scored according to the following metric: sign(*logFC between ARID5B-deficient and control*) x -log_10_(*p-value*). Where multiple peaks corresponded to a single gene, genes were ranked according to the peak with the largest absolute value score. We used fgsea (v1.20.0) to perform gene set enrichment analysis on this ranked gene set. We used trackplot^75^ to visualize ATAC-seq peaks.

#### Single-cell RNA sequencing analysis

Single-cell RNA sequencing read counts and metadata are publicly available for the AMP RA/SLE Phase 2 synovial tissue dataset (Synapse, accession 52297480; https://www.synapse.org/Synapse:syn52297840.7/datasets/)^9^. Cell projections in UMAP space from the original study were used for visualization and gene expression plotting. Seurat (v4.3.0)^79^ was used for data processing, normalization, and differential expression analysis.

Read counts for each cell were normalized by multiplicatively scaling to 10,000 reads followed by log normalization (NormalizeData function). Inflammatory and invasive gene expression program scores for each cell were computed using the AddModuleScore function based on the gene lists shown in Table S1. Fibroblasts scoring in the top tenth percentile for either gene signature were designated as inflammatory or invasive. Genes differentially expressed between inflammatory and invasive fibroblasts were identified using the Wilcoxon rank sum test with Bonferroni p-value correction. Transcription factors were scored and ranked according to the following metric: sign(*logFC between inflammatory and invasive*) x -log_10_(*adjusted p-value*) x (*proportion expression in the upregulated fibroblast population*). Gene expression program scores for ARID5B-upregulated and ARID5B-downregulated gene signatures were computed using the AddModuleScore function.

#### K/BxN serum transfer-induced arthritis

Arthritogenic K/BxN serum was purchased from Dr. Peter Nigrovic, Division of Immunology, Boston Children’s Hospital, Boston, MA. K/BxN serum transfer-induced arthritis was induced by two intravenous (retro-orbital) injections of serum from KRN mice: (1) 100µl serum on day 0, and (2) an additional 50µl serum on day 2. Forepaw and hindpaw joint swelling was assessed daily, both objectively (by caliper measurement) and subjectively (by clinical scoring). For clinical scoring, each of four paws was scored on a scale from 0-4 with a maximal score of 16. Normal paws received a score of 0; one point was given for swelling involving each aspect of the joint (digits, ventral surface, dorsal surface, wrist/ankle joint).

Unless otherwise indicated, 6-12-week-old male mice and littermate controls were used for all arthritis experiments, as female mice do not respond as efficiently or uniformly in this disease model^80^. Data shown represent pooled results from 2-3 independent experiments. Researchers were blinded throughout the collection of clinical and caliper measurements.

#### Mouse synovial tissue histology

Paws were harvested at days 8 and 14 of STIA and were fixed in paraformaldehyde (4%), decalcified in EDTA (10% w/v, pH 7.4) for 3 weeks, and embedded in paraffin. 7µm sagittal sections were cut and stained with either hematoxylin/eosin (H&E, Abcam) or with safranin O (SafO, Sigma) and Fast Green (Sigma) per manufacturer instructions. At least four H&E-stained sections and four SafO/Fast Green-stained sections per specimen (each set spanning a total range of least 70µm) were used for histopathologic assessment and scoring based on Hayer et al.^81^ with modifications as described below.

Synovial inflammation was scored on a scale from 0-4 as follows: 0 – normal synovial membrane (1-2 cells thickness) with no inflammatory infiltrate; 1 – mildly hyperplastic synovial membrane (3-5 cells thickness) with sparse inflammatory infiltrate; 2 – hyperplastic synovial membrane (≥5 cells thickness) with enhanced inflammatory infiltrate in some but not all joints; 3 – hyperplastic synovial membrane with enhanced inflammatory infiltrate in most joints; 4 – hyperplastic synovial membrane with diffuse inflammatory infiltrate involving entire joints and surrounding connective tissue.

Bone and cartilage erosion were scored on a scale from 0-4 as follows: 0 – healthy, intact bone with smooth, fully SafO-stained articular cartilage layers; 1 – small, superficial erosions of the outer cortical bone and partial SafO destaining or loss of up to one-third of the superficial cartilage layer; 2 – enhanced erosions penetrating most of the cortical bone and SafO destaining or loss of up to one-half of the superficial cartilage layer; 3 – enhanced bone erosions penetrating the cortical layer with focal breakthrough to marrow cavity and diffuse SafO destaining or thinning of the superficial cartilage layer; 4 – massive bone erosions penetrating the cortical layer with significant breakthrough to marrow cavity and complete SafO destaining or loss of the superficial cartilage layer.

Inflammation and erosion were scored for each joint within a given section, and mean scores per section were averaged to compute a composite histologic inflammation score and a histologic erosion score for each specimen. Scores were then averaged across each experimental group.

