## Supplemental Data for "ARID5B drives an inflammatory-to-destructive shift in pathologic fibroblast behavior"

**Supplemental information**

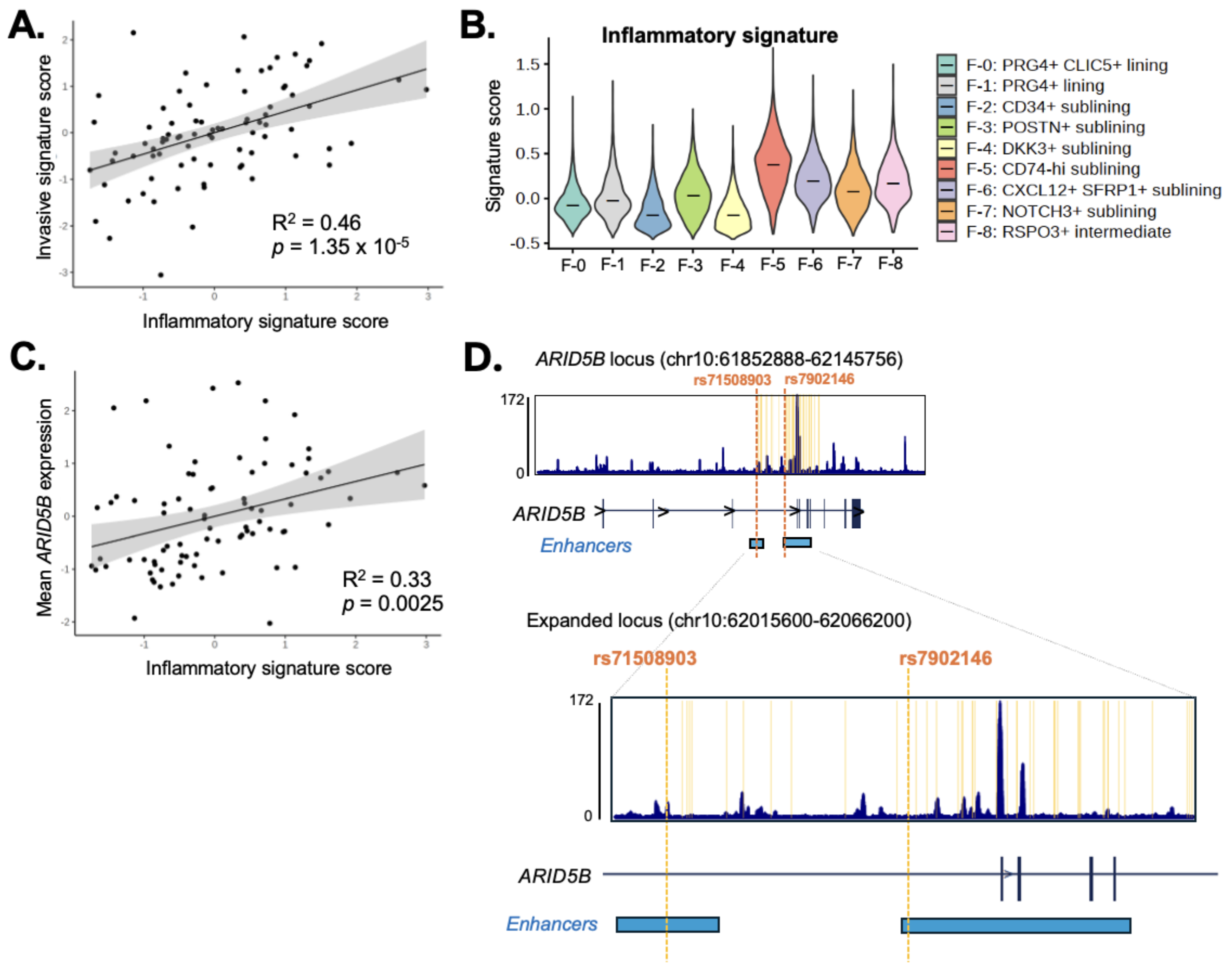


**Figure S1. *ARID5B* expression across fibroblast populations and RA patients, related to Figure 1. (A)** Scatterplot of scaled mean fibroblast inflammatory and invasive gene signature enrichment scores for each patient from the AMP RA/SLE Phase 2 study[9]. $R^2$ and p-value from Pearson correlation test. **(B)** Violin plots depicting inflammatory gene signature enrichment scores across the indicated synovial fibroblast subsets. Median expression indicated. **(C)** Scatterplot of scaled mean fibroblast inflammatory gene signature enrichment scores and *ARID5B* expression levels for each patient from the AMP RA/SLE Phase 2 study. $R^2$ and p-value from Pearson correlation test. **(D)** Track plot depicting ATAC-seq peaks along the *ARID5B* gene locus in synovial fibroblasts (averaged signals from n = 2 primary cell lines). Orange lines indicate genomic coordinates of SNPs significantly associated with RA risk ($p < 1 \times 10^{-8}$ in Ishigaki et al.[43]; lead SNPs labeled), shown in relation to the fibroblast ATAC-seq peaks and predicted *ARID5B* gene enhancer elements.

### Table S1. Genes included in the inflammatory and invasive gene modules used in single-cell transcriptomic analysis of RA synovial fibroblasts, related to Figure 1.

| Gene symbol | Category | Description | References (PMID) |
|---|---|---|---|
| IL6 | Inflammatory | Pleiotropic cytokine with roles in T/B cell differentiation and activation and acute phase response, RA synovial fibroblasts are major source | 32327746, 31061532 |
| LIF | Inflammatory | IL-6 family member cytokine that amplifies RA synovial fibroblast inflammatory factor production via an autocrine signaling loop | 1522240, 28228280 |
| CXCL8 | Inflammatory | Neutrophil chemoattractant and proangiogenic factor highly expressed by RA synovial fibroblasts | 7561066, 11673556, 36725964 |
| CXCL12 (SDF1) | Inflammatory | Monocyte and lymphocyte chemoattractant highly expressed by RA synovial fibroblasts | 11086103, 11920421, 36725964 |
| CCL2 (MCP1) | Inflammatory | Monocyte and lymphocyte chemoattractant, highly expressed by RA synovial fibroblasts and further stimulates their inflammatory factor production | 33453247, 11673556 |
| TNFSF13B (BAFF) | Inflammatory | B cell survival and maturation factor implicated in supporting autoreactive B cell activation and class-switching | 21798884, 15634908, 19737141 |
| HLA-DRA | Inflammatory | MHC class II molecule highly expressed by RA synovial fibroblasts, involved in antigen presentation to autoreactive T/B cells | 28649674, 31061532 |
| CD74 | Inflammatory | Invariant chain; facilitates MHC class II molecule assembly and cell surface expression, highly expressed by RA synovial fibroblasts | 37938773, 40093895 |
| CCL19 | Inflammatory | Chemokine for T cell recruitment and activation, highly expressed by RA synovial fibroblasts | 21225692, 34651581 |
| CCL20 | Inflammatory | Chemokine for monocyte/T cell recruitment and activation, highly expressed by RA synovial fibroblasts | 19447772, 24394994 |
| CXCL1 | Inflammatory | Neutrophil chemoattractant, highly expressed by RA synovial fibroblasts and further stimulates their inflammatory factor production | 7561066, 33087182 |
| CXCL3 | Inflammatory | Chemokine for monocyte recruitment and angiogenesis, highly expressed by RA synovial fibroblasts | 32079724, 36860872 |
| CXCL6 | Inflammatory | Chemokine for neutrophil recruitment, highly expressed by RA synovial fibroblasts | 32079724, 20036936 |
| VCAM1 | Inflammatory | Binds leukocyte integrins to mediate leukocyte migration and tissue infiltration, highly expressed by RA synovial fibroblasts | 8909250, 10820282 |
| STAT1 | Inflammatory | JAK/STAT transcription factor, significantly induced in cytokine-stimulated RA synovial fibroblasts | 37938773, 14962955 |
| CDH11 | Invasive/Remodeling | Mesenchymal cadherin involved in synovial lining formation and RA synovial fibroblast-mediated joint destruction | 17255475, 19404963, 22127696 |
| MMP1 | Invasive/Remodeling | Collagenase targeting intact fibrillar collagens, highly expressed by RA synovial fibroblasts | 11714387, 37938773 |
| MMP2 | Invasive/Remodeling | Gelatinase degrading collagen II, highly expressed at the invasive pannus surface by synovial fibroblasts | 9923652, 18471998 |
| MMP3 | Invasive/Remodeling | Aggrecanase; levels correlate with erosive damage in human RA, highly expressed by RA synovial fibroblasts | 10556259, 37938773 |
| MMP9 | Invasive/Remodeling | Gelatinase degrading collagen monomers, enhances RA synovial fibroblast migration and invasion | 14532148, 22960198 |
| MMP13 | Invasive/Remodeling | Collagenase targeting intact collagen II/fibrillar collagens and implicated in cartilage destruction, highly expressed by RA synovial fibroblasts | 11714387, 18289056 |
| MMP14 | Invasive/Remodeling | Membrane-anchored proteolytic enzyme that activates other MMPs to facilitate ECM degradation, implicated in pannus invasion in RA synovium | 9923652, 19248098 |
| CD44 | Invasive/Remodeling | Adhesion molecule that binds hyaluronan; highly expressed by RA synovial fibroblasts and modulates their migratory and invasive behavior | 10943861, 15781582 |
| PDPN | Invasive/Remodeling | Transmembrane glycoprotein promoting RA synovial fibroblast-mediated cartilage and bone destruction, possibly through invadopodia regulation | 27863512, 25486435 |
| TNFSF11 (RANKL) | Invasive/Remodeling | Cytokine required for osteoclastogenesis and bone erosion; RA synovial fibroblasts are major source | 26025971, 16490750 |
| EZR | Invasive/Remodeling | Crosslinker between actin cytoskeleton and plasma membrane molecules including CD44; promotes cell migration and invasion | 24599913, 12711360 |
| EGFR | Invasive/Remodeling | Growth factor receptor highly expressed on RA synovial fibroblasts, implicated in promoting invasiveness and osteoclast activation | 31068444, 9683636, 22393153 |
| HBEGF | Invasive/Remodeling | Growth factor highly expressed on RA synovial fibroblasts, implicated in promoting fibroblast motility and invasiveness | 31068444, 9683636 |
| PDGFRA | Invasive/Remodeling | Growth factor receptor highly expressed on RA synovial fibroblasts, implicated in promoting fibroblast proliferation, motility and invasiveness | 1724096, 26704941 |
| PDGFRB | Invasive/Remodeling | Growth factor receptor highly expressed on RA synovial fibroblasts, implicated in promoting fibroblast proliferation, motility and invasiveness | 1724096, 26704941 |

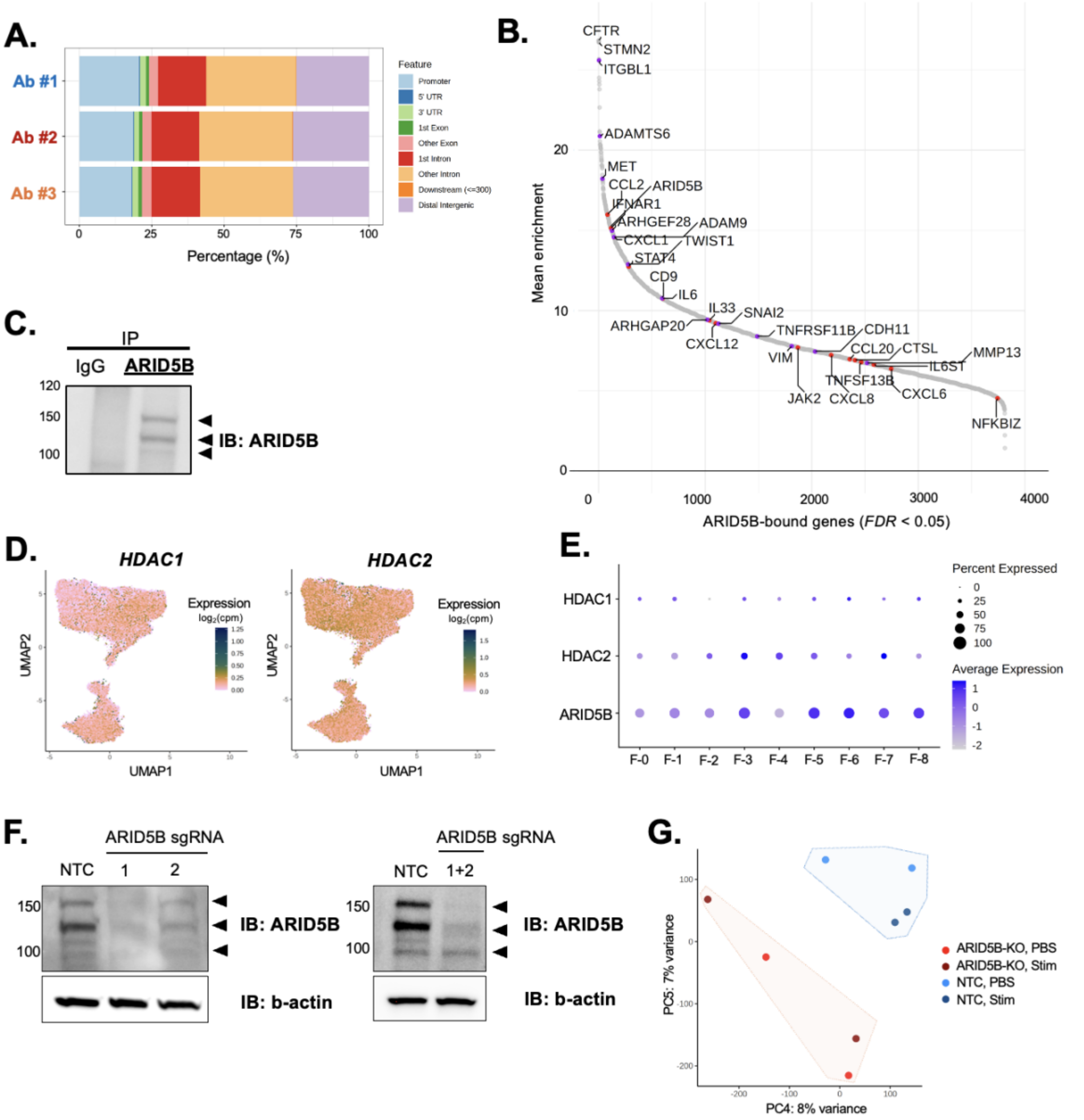


**Figure S2. ARID5B genomic binding and epigenetic regulation, related to Figure 2. (A)** Distribution of ARID5B binding sites in relation to annotated genomic features in synovial fibroblasts. Each row corresponds to a single CUT&RUN sequencing experiment performed using the indicated anti-ARID5B antibody. **(B)** Plot of mean peak enrichment values for genes with regulatory loci bound by ARID5B in all three ARID5B CUT&RUN sequencing reactions *(FDR* < 0.05)*.* Purple points indicate genes associated with cell motility and invasion pathways; red points indicate genes associated with inflammation and immune regulation pathways. **(C)** Immunoblot for ARID5B following ARID5B immunoprecipitation from synovial fibroblast lysates; pulldown with nonspecific IgG shown as negative control. ARID5B isoforms indicated by arrows. **(D)** Projection of *HDAC1* and *HDAC2* expression levels across synovial fibroblasts in UMAP space. **(E)** Dot plot of scaled mean and proportion of *HDAC1*, *HDAC2*, and *ARID5B* expression in each of the indicated synovial fibroblast subsets. **(F)** Immunoblot for ARID5B following CRISPR/Cas9-mediated deletion of ARID5B with the sgRNAs listed compared to wild-type non-targeting control sgRNA (NTC). ARID5B isoforms indicated by arrows. Beta-actin shown as cell lysate loading control. **(G)** PCA of normalized and scaled peak counts in wild-type (NTC) and ARID5B-deficient (KO) fibroblasts. Samples are colored by genotype and stimulation status.

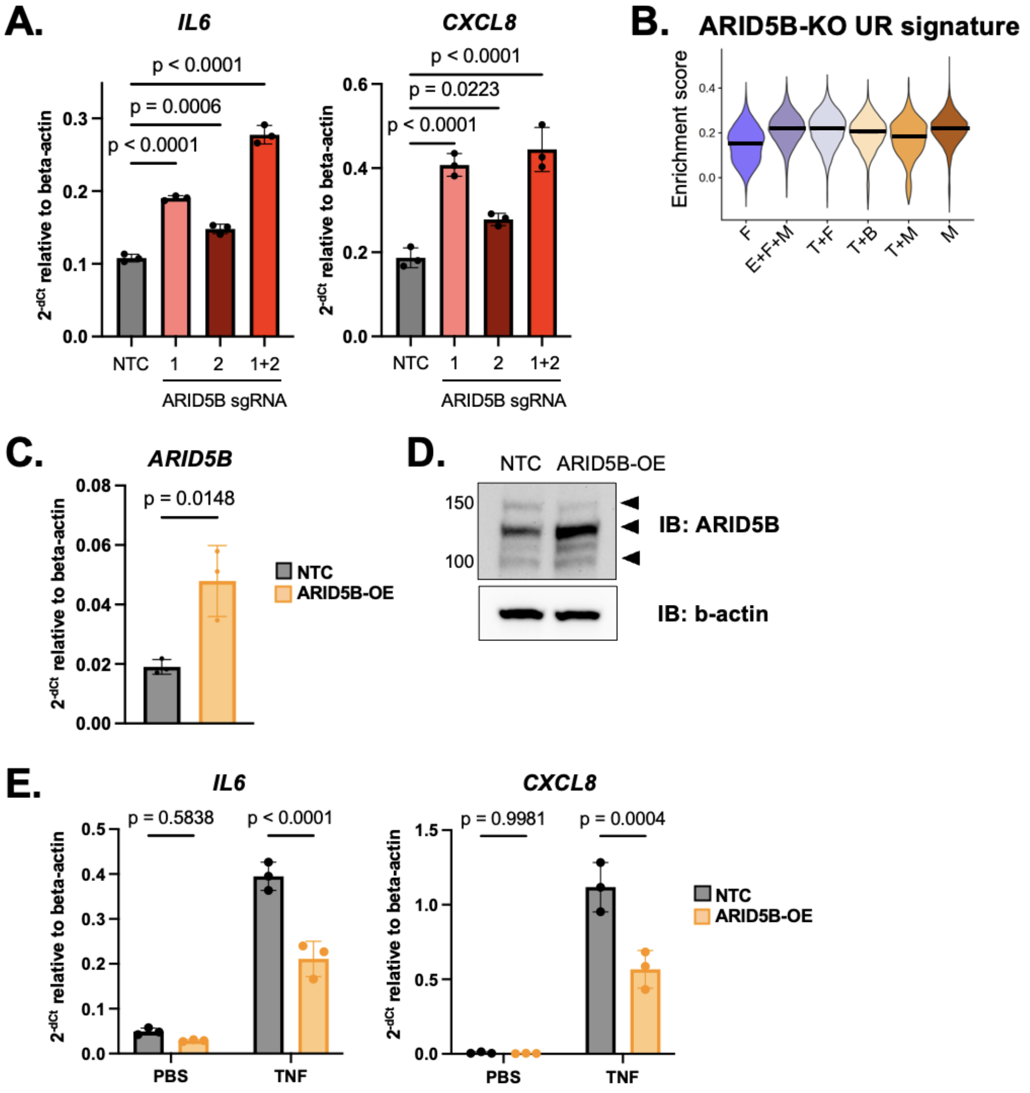


**Figure S3. Inflammatory factor suppression by ARID5B, related to Figure 3. (A)** RT-qPCR of *IL6* and *CXCL8* gene expression in TNFα-stimulated control (NTC) and ARID5B-deficient synovial fibroblasts. Mean ± SD shown. P values from one-way ANOVA with Dunnett multiple comparisons test. **(B)** Violin plot of ARID5B-upregulated gene signature scores across fibroblasts from all RA CTAPs. Median enrichment values shown. **(C)** RT-qPCR of *ARID5B* expression in fibroblasts following CRISPRa-mediated overexpression of *ARID5B* (OE) as compared to control (NTC). Mean ± SD shown. P value from two-sided Student's t test. **(D)** Immunoblot for ARID5B in ARID5B-overexpressing fibroblasts compared to control fibroblasts (NTC). ARID5B isoforms indicated by arrows. Beta-actin shown as cell lysate loading control. **(E)** RT-qPCR of *IL6* and *CXCL8* gene expression in unstimulated (PBS) and TNFα-stimulated control (NTC) and ARID5B-overexpressing (OE) synovial fibroblasts. Mean ± SD shown. P values from two-way ANOVA with Šídák's multiple comparisons test.

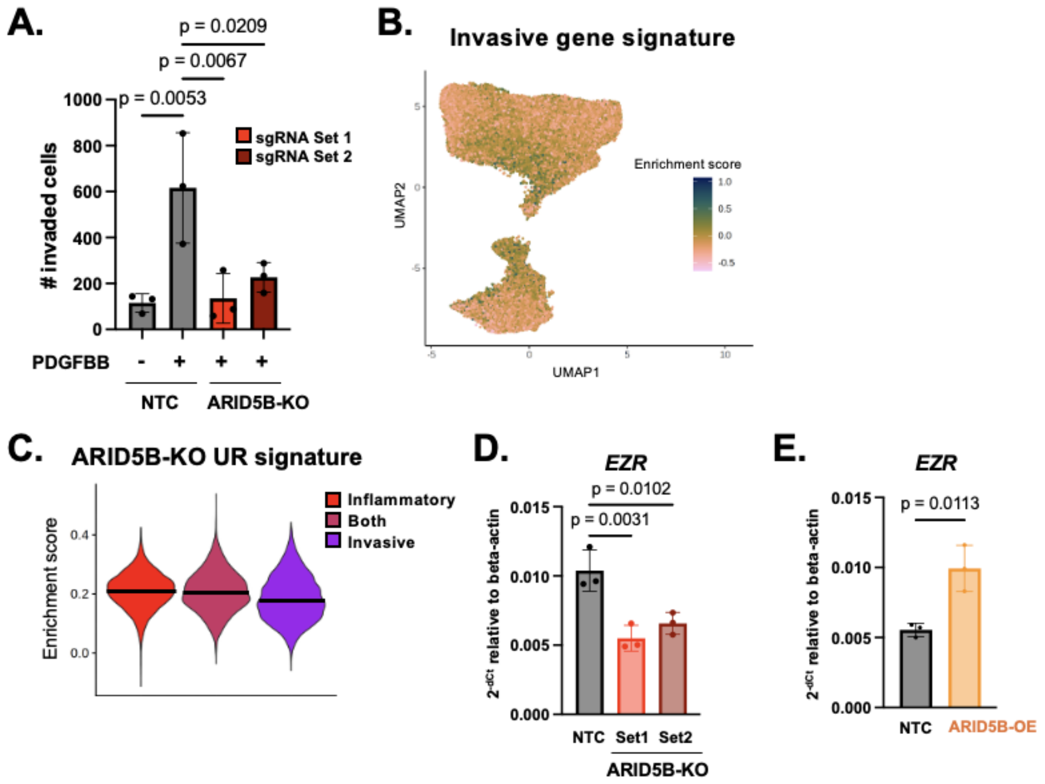


**Figure S4. Upregulation of fibroblast invasion and *EZR* expression by ARID5B, related to Figure 4. (A)** Enumeration of invaded cells by control (NTC) and ARID5B-deficient (KO) fibroblasts. Mean ± SD shown. P values from one-way ANOVA with Dunnett multiple comparisons test. **(B)** Projection of invasive gene signature enrichment scores across synovial fibroblasts in UMAP space **(C)** Violin plots of enrichment scores for the gene signature upregulated by ARID5B deletion in inflammatory versus invasive and mixed inflammatory-invasive synovial fibroblasts. **(D)** RT-qPCR of *EZR* gene expression in control (NTC) and ARID5B-deficient (KO) fibroblasts. Mean ± SD shown. P values from one-way ANOVA with Dunnett multiple comparisons test. **(E)** RT-qPCR of *EZR* gene expression in control (NTC) and ARID5B-overexpressing (OE) fibroblasts. Mean ± SD shown. P values from two-sided Student's t test.

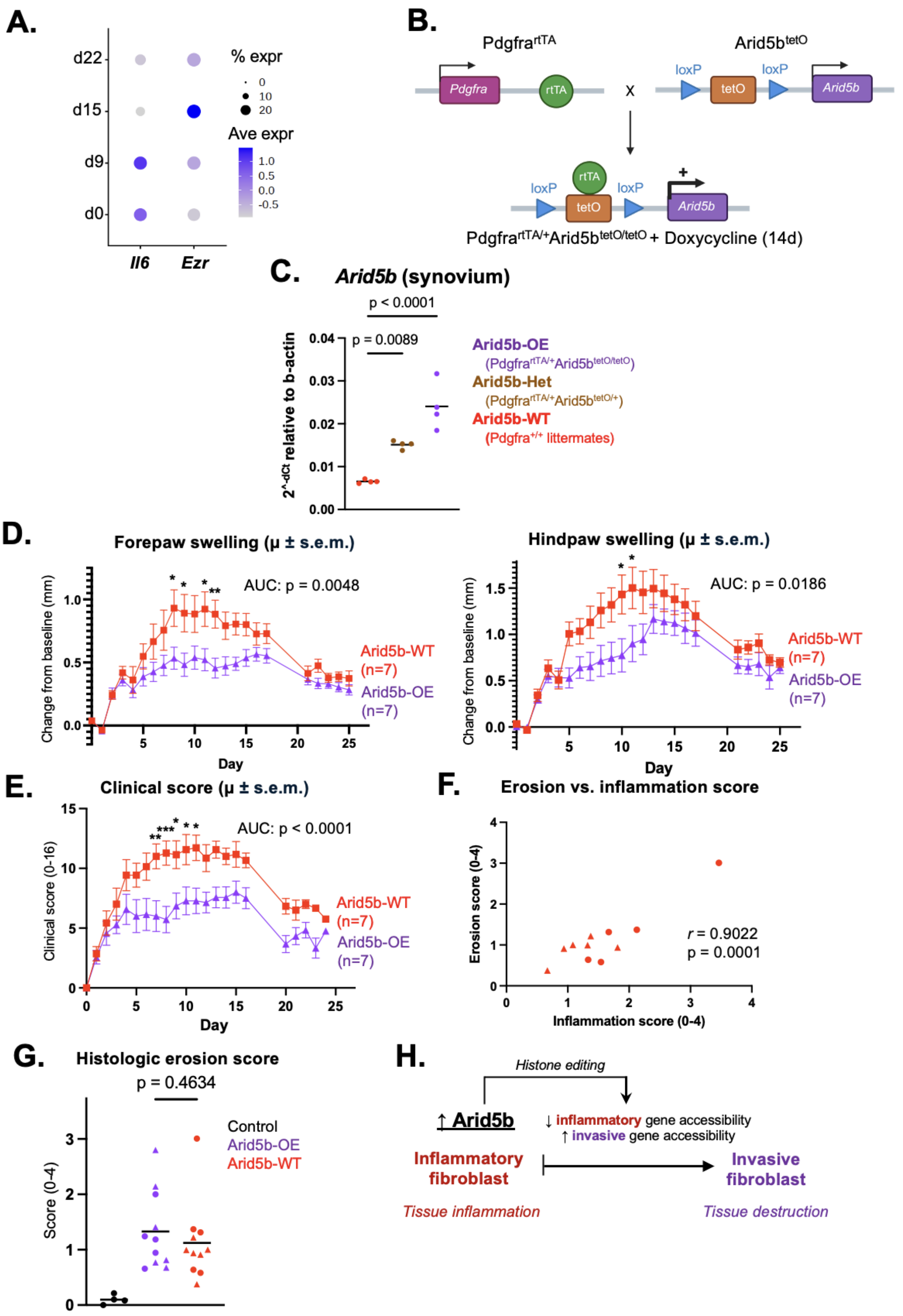

A.
% expr
Ave expr
d22
d15
d9
d0
Il6
Ezr
B.
PdgfrartTA
Arid5btetO
Pdgfra
rtTA
loxP
tetO
Arid5b
PdgfrartTA/+Arid5btetO/tetO + Doxycycline (14d)
C.
Arid5b (synovium)
p < 0.0001
p = 0.0089
2^-dCt relative to b-actin
Arid5b-OE
(PdgfrartTA/+Arid5btetO/tetO)
Arid5b-Het
(PdgfrartTA/+Arid5btetO/+)
Arid5b-WT
(Pdgfra+/+ littermates)
D.
Forepaw swelling (μ ± s.e.m.)
AUC: p = 0.0048
Change from baseline (mm)
Day
Arid5b-WT (n=7)
Arid5b-OE (n=7)
Hindpaw swelling (μ ± s.e.m.)
AUC: p = 0.0186
E.
Clinical score (μ ± s.e.m.)
AUC: p < 0.0001
Clinical score (0-16)
F.
Erosion vs. inflammation score
Erosion score (0-4)
Inflammation score (0-4)
r = 0.9022
p = 0.0001
G.
Histologic erosion score
p = 0.4634
Score (0-4)
Control
Arid5b-OE
Arid5b-WT
H.
Histone editing
↑ Arid5b
↓ inflammatory gene accessibility
↑ invasive gene accessibility
Inflammatory fibroblast
Invasive fibroblast
Tissue inflammation
Tissue destruction

**Figure S5. Assessment of arthritic disease in mice with fibroblast-specific overexpression of Arid5b, related to Figure 5. (A)** Dot plot of scaled mean and proportion of *Il6* and *Ezr* expression in synovial fibroblasts at each of the indicated STIA timepoints. **(B)** Schematic describing the generation of Arid5b-OE (Pdgfra$^{rtTA/+}$Arid5b$^{tetO/tetO}$) mice. **(D)** RT-qPCR of *Arid5b* gene expression in knee synovium harvested from Arid5b-OE, Arid5b-Het, and Arid5b-WT mice following two weeks of doxycycline administration. P values by one-way ANOVA. **(D-E)** Caliper measurements **(D)** and clinical scoring **(E)** of paw swelling and inflammation in Arid5b-OE mice (n = 7) and Arid5b-WT littermates (n = 7). Mean ± SEM shown at each timepoint. *p < 0.05; **p < 0.01; ***p < 0.001; ****p < 0.0001 by two-way repeated measures ANOVA with Šídák's multiple comparisons test. **(F)** Scatterplot of histologic erosion versus inflammation scores in Arid5b-WT mice (circles: n = 5 from STIA day 8; triangles: n = 6 from STIA day 14). *R* and p values by two-tailed Pearson's correlation test. **(G)** Histologic erosion scores in nonarthritic control (n = 4) and arthritic Arid5b-OE and Arid5b-WT mice (circles: n = 5 from STIA day 8; triangles: n = 6 from STIA day 14 per group). P value from two-sided Student's t test. **(H)** Model for Arid5b function in pathologic tissue fibroblasts.
